# Voltage-gated calcium channels contribute to spontaneous glutamate release directly via nanodomain coupling or indirectly via calmodulin

**DOI:** 10.1101/2020.11.10.376111

**Authors:** Byoung Ju Lee, Che Ho Yang, Seung Yeon Lee, Suk-Ho Lee, Yujin Kim, Won-Kyung Ho

## Abstract

Neurotransmitter release occurs either synchronously with action potentials (evoked release) or spontaneously (spontaneous release). Whether the molecular mechanisms underlying evoked and spontaneous release are identical, especially whether voltage-gated Ca^2+^ channels (VGCCs) can trigger spontaneous events, is still a matter of debate in glutamatergic synapses. To elucidate this issue, we characterized the VGCC dependence of miniature excitatory postsynaptic currents (mEPSCs) in various synapses with different coupling distances between VGCCs and synaptic vesicles, known as a critical factor in evoked release. We found that most of the extracellular calcium-dependent mEPSCs were attributable to VGCCs in cultured autaptic hippocampal neurons and the mature calyx of Held where VGCCs and vesicles were tightly coupled. Among loosely coupled synapses, mEPSCs were not VGCC-dependent at immature calyx of Held and CA1 pyramidal neuron synapses, whereas VGCCs contribution was significant at CA3 pyramidal neuron synapses. Interestingly, the contribution of VGCCs to spontaneous glutamate release in CA3 pyramidal neurons was abolished by a calmodulin antagonist, calmidazolium. These data suggest that coupling distance between VGCCs and vesicles determines VGCC dependence of spontaneous release at tightly coupled synapses, yet VGCC contribution can be achieved indirectly at loosely coupled synapses.

**Highlights:**

- Extracellular Ca^2+^ -dependent spontaneous glutamate release is mediated by VGCCs.
- Ca^2+^ cooperativity of spontaneous and evoked release is comparable.
- Synapses at different areas and ages show distinct VGCC dependence on mini events.
- Nanodomain coupling between VGCCs and Ca^2+^ sensors determines VGCC dependence.

## INTRODUCTION

Synaptic transmission is a major mechanism of information processing in neural systems. Neuronal excitation can be transmitted to postsynaptic neurons via neurotransmitter release triggered by action potentials (APs) in presynaptic nerve terminals. However, neurotransmitter release also occurs spontaneously at resting state with a low frequency of around 0.01 Hz per synaptic bouton (Murthy and Stevens, 1999; Sara et al., 2005). This type of synaptic communication plays a pivotal role in synaptic structure and function, including synapse maturation and maintenance, homeostasis, and plasticity (Kavalali, 2015). Despite the physiological importance of spontaneous release, its molecular mechanisms are poorly understood.

Neurotransmitter release occurs when an increase in intracellular Ca^2+^ is detected by Ca^2+^ sensors at presynaptic terminals. During AP firing, Ca^2+^ influx mediated by voltage-gated Ca^2+^ channels (VGCCs) is captured by vesicular Ca^2+^ sensors, leading to synaptic vesicle fusion. Spontaneous release also involves Ca^2+^-dependent processes, yet it is not clear whether Ca^2+^-dependent spontaneous release utilizes the same Ca^2+^ sensors and sources as the evoked release (Groffen et al., 2010; Kavalali, 2015; Xu et al., 2007; Xu et al., 2009). In particular, the contribution of VGCCs to spontaneous release remains controversial. It is generally accepted that stochastic VGCC activity is a major trigger of spontaneous release at inhibitory synapses (Goswami et al., 2012; Williams et al., 2012), yet many studies deny a contribution of VGCCs at excitatory synapses (Courtney et al., 2018; Dai et al., 2015; Tsintsadze et al., 2017; Vyleta and Smith, 2011). It has been suggested that glutamatergic spontaneous release is independent of Ca^2+^ influx via VGCCs, but tonically activated by the calcium-sensing receptor (CaSR), which is a G-protein coupled receptor whose activity depends on external Ca^2+^ concentration ([Ca^2+^]e) (Tsintsadze et al., 2017; Vyleta and Smith, 2011). On the other hand, Ermolyuk et al. (2013) showed clear evidence that glutamatergic spontaneous release in cultured hippocampal neurons is triggered by local Ca^2+^ increases induced by stochastic opening of presynaptic VGCCs at the resting state. These contradictory results may imply that spontaneous release is mediated by multiple Ca^2+^ sensors and multiple Ca^2+^ sources, and their relative contributions are distinct among different cell types and differentially regulated by various cellular states or signaling mechanisms.

The coupling distance between Ca^2+^ sources and Ca^2+^ sensors of synaptic vesicles is a key determinant of the efficacy and speed of AP-triggered synaptic transmission (Eggermann et al., 2012; Neher and Sakaba, 2008). In nanodomain coupling, a single or a few Ca^2+^ channel openings would lead to brief and local increases in Ca^2+^ concentration and directly trigger synaptic vesicle fusion, so that tight coupling has functional advantages, in terms of speed, temporal precision, and energy efficiency of synaptic transmission (Eggermann et al., 2012; Schmidt et al., 2013). On the other hand, loose coupling enables the control of initial release probability by fast endogenous Ca^2+^ buffers and the generation of facilitation by buffer saturation, providing the molecular framework for presynaptic plasticity (Vyleta and Jonas, 2014). The coupling configuration may also have implications for spontaneous release, but little information is available to date.

In the present study, we investigated the contribution of VGCCs to spontaneous release at various types of glutamatergic synapses in the brain. We found that Ca^2+^-dependent spontaneous release is mediated mostly by VGCCs at tightly coupled synapses on cultured autaptic hippocampal neurons and the mature calyx of Held, whereas VGCCs contribute little to spontaneous release from loosely coupled synapses on immature calyx of Held and CA1 pyramidal neurons (CA1-PCs), supporting the idea that the coupling distance between Ca^2+^ sensors and sources is the key determinant of VGCC dependence of spontaneous release. However, spontaneous release from loosely coupled synapses on CA3 pyramidal neurons (CA3-PCs) was also VGCC-dependent, which was abolished in the presence of calmidazolium, an inhibitor of calmodulin. These data suggest distinct regulatory mechanisms of spontaneous release in various types of neurons.

## RESULTS

### Multiple Ca^2+^ channels contribute to spontaneous glutamate release in autaptic hippocampal neurons

To measure spontaneous release at excitatory synapses, we recorded synaptic activities from isolated hippocampal neurons grown on astrocyte feeder islands for at least 3 weeks for synaptic maturation (Fig. 1Aa). Recordings were performed at resting state (−70 mV) under whole-cell voltage-clamp conditions using pipette solutions containing 0.1 mM ethylene glycol-bis (β-aminoethyl ether)- N,N,N′,N′-tetraacetic acid (EGTA). The cells were first identified as glutamatergic neurons based on the fast decay kinetics of synaptic currents (decay time constant: 3.02 ± 0.16 ms, N = 64), and their identity was subsequently confirmed pharmacologically. Postsynaptic currents that decayed within 10 ms were not affected by 0.1 mM picrotoxin but were abolished by the AMPA receptor antagonist, CNQX (10 μM) (Fig. 1Ab). These activities were not affected by tetrodotoxin (Fig. 1Ac), confirming that none of the synaptic events were activity-dependent in voltage-clamped cultured autaptic neurons. Therefore, we regarded these events as miniature excitatory postsynaptic currents (mEPSCs) representing spontaneous glutamate release. Frequencies of mEPSCs were variable among cells, ranging from 0.5 to 10 Hz (Fig. 1Ab. Bottom right, mean = 3.87 ± 0.18 Hz, N = 278). Hereafter, the mEPSC frequencies obtained after applying any experimental conditions were normalized to the control level obtained in normal ACSF containing 2 mM Ca^2+^. After the removal of external Ca^2+^ (nominally Ca^2+^-free), the mEPSC frequency decreased to 0.42 ± 0.03 (N = 13) (Figs. 1Ba, Bd), whereas inhibition of internal Ca^2+^ release from the endoplasmic reticulum (ER) via ryanodine receptors (RyRs) using ryanodine (10 μM) reduced the mEPSC frequency to 0.85 ± 0.04 (N = 6) (Figs. 1Bb, Bd). Ryanodine still reduced the mEPSC frequency in Ca^2+^-free conditions (Supplementary Fig. 1), suggesting that internal and external Ca^2+^ sources mediate spontaneous release through independent mechanisms. Inhibition of IP3 receptor (IP3R)-induced Ca^2+^ release from the ER by 2-aminoethoxydiphenylborane (2-APB, 10 μM) reduced the mEPSC frequency (0.82 ± 0.04, N = 9, P = 0.0002) to a level comparable to that in the presence of ryanodine (Figs. 1Bc, Bd). There was no additive effect when both ryanodine and 2-APB were applied (Figs. 1Bb to Bd), suggesting that RyRs and IP3Rs share internal Ca^2+^ stores. Taken together, 27% and 15% of miniature events in cultured autaptic hippocampal neurons were attributable to Ca^2+^-independent and ER-dependent mechanisms, respectively, while 58% were [Ca^2+^]e-dependent (Fig. 1Be). The mEPSC frequency changed according to [Ca^2+^]e changes (Fig. 1C), with the slope of the log-log plot for mEPSC frequency against [Ca^2+^]e being 0.42 (black symbols and line, Fig. 1D), which is compatible with the value measured in cortical neurons (0.63, (Vyleta and Smith, 2011)) However, Ca^2+^ cooperativity shown in this slope is underestimated, as 42% of all mEPSCs are independent of [Ca^2+^]e. To assess the actual Ca^2+^ cooperativity, we subtracted the [Ca^2+^]e-independent fraction and recalculated the slope. The result was 1.24 (red line, Fig. 1D), which was comparable to the Ca^2+^ cooperativity of evoked excitatory postsynaptic currents (eEPSCs) obtained in hippocampal synapses (1.77: Vyleta and Jonas, 2014), suggesting that mechanisms underlying [Ca^2+^]e-dependent spontaneous release may not be substantially different from those of evoked release.

**Fig. 1.**
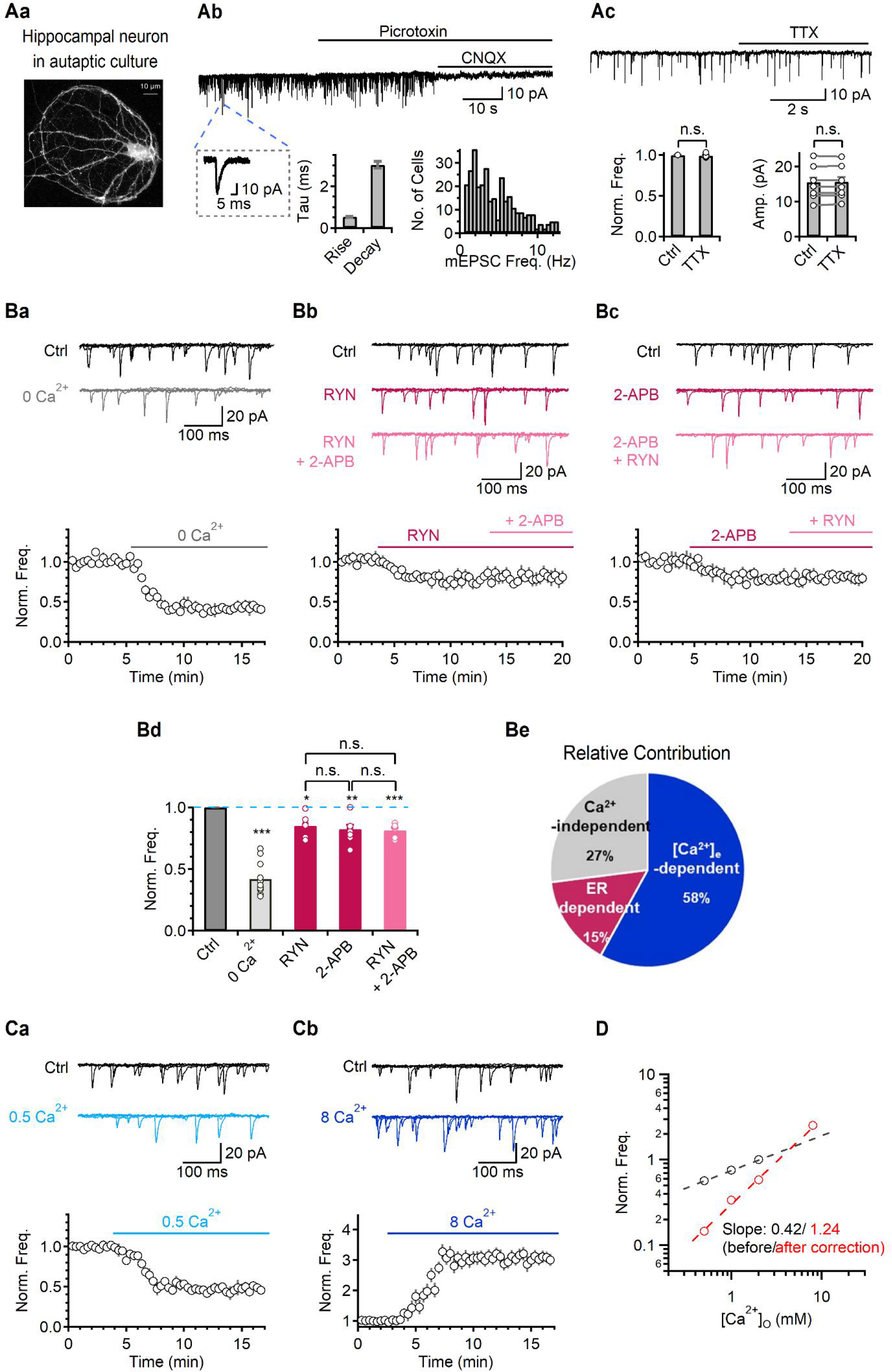
Ca^2+^ cooperativity for glutamatergic spontaneous release. (Aa) A representative image of hippocampal neurons in autaptic culture system labeled by the extracellular application of lipophilic fluorescent dyes, Dil. (Ab) Top. A representative trace of mEPSC in the presence of picrotoxin (PTX) and CNQX. Bottom. left: An enlarged trace of a single mEPSC. middle: A bar graph indicating rise and decay time constants of mEPSCs. Rise time constant was 0.53 ± 0.02 ms (*N* = 62). right: A histogram representing number of cells by mEPSC frequency. (Ac) A representative trace of mEPSCs before (Ctrl) and after the application of 0.5 μM tetrodotoxin (TTX). (B) Ba-Bc. Top. Representative traces of mEPSCs in control (2 mM Ca^2+^) and 0 mM [Ca^2+^]_e_ (0 Ca^2+^) (Ba), and the presence of ryanodine (RYN) followed by 2-aminoethoxydiphenylborane (2-APB) (Bb), and 2-APB followed by RYN (Bc), respectively. Five 500 ms-long mEPSC traces were overlaid. Bottom. Average time courses of the normalized mEPSC frequency. In each time course plot, solid lines indicate the presence of 0 Ca^2+^, RYN, 2-APB, and RYN with 2-APB, respectively. The data were normalized by the mean mEPSC frequency of control. (Bd) A bar graph of average values of the normalized mEPSC frequency in 0 Ca^2+^ (*N* = 13, 0.42 ± 0.03, *P*<0.001; Ba), RYN (*N* = 6, 0.85 ± 0.04; Bb), 2-APB (*N* = 9, 0.82 ± 0.04; Bc) or RYN with 2-APB (*N* = 6, 0.81 ± 0.02; Bb). (Be) A pie chart evaluating the relative contribution of [Ca^2+^]_e_- dependent (blue), Ca^2+^-independent (grey) or ER-dependent (magenta) mEPSCs to the total mEPSCs. (Ca, Cb) Top. Representative traces of mEPSCs in different [Ca^2+^]_e_ from 2 (Ctrl) to 0.5 mM (0.5 Ca^2+^, light blue) (Ca) and from 2 to 8 mM (8 Ca^2+^, dark blue) (Cb). Five 500 ms-long mEPSC traces were overlaid. External [Mg^2+^] was 1 mM for these recordings. Bottom. An average time course of the normalized mEPSC frequency (Ca, 0.5 Ca^2+^, 0.56 ± 0.02, N = 28; Cb, 8 Ca^2+^, 2.94 ± 0.24, N = 17; compared to control). In each time course plot, solid lines indicate the presence of 0.5 and 8 Ca^2^, respectively. The data were normalized by the mean mEPSC frequency of control. (D) A log-log plot for the normalized mEPSC frequency against [Ca^2+^]_e_. The slope of mEPSC frequency was 0.42 which was fitted from control (2 mM Ca^2+^) to lower concentration (black circles and dash line). To overcome underestimation, the slope was recalculated after the 0 Ca^2+^ fraction was subtracted (red circles and dashed line, Slope; 1.24). All data are represented as mean ± S.E.M., \**P*<0.05, \*\*\**P*<0.001, single group mean *t*-test or paired *t*-test (Bd); n.s. = not significant.

### VGCCs contribute to spontaneous glutamate release

It is well known that evoked release is triggered by Ca^2+^ influx through presynaptic VGCCs during action potentials, but the contribution of presynaptic VGCCs to spontaneous glutamate release remains controversial (Ermolyuk et al., 2013; Vyleta and Smith, 2011). To test this possibility, we examined the contribution of VGCCs to mEPSCs using blockers to specific VGCC subtypes. Inhibition of P/Q-type (with 0.1 μM ω-agatoxin-IVA [Aga]), N-type (with 0.1 μM ω-conotoxin GVIA [Cono]), and R-type (with 0.3 μM SNX-482 [SNX] or 100 μM NiCl2) VGCCs significantly decreased the mEPSC frequency (Figs. 2Aa-c, C; Aga, 0.71 ± 0.02, N = 24; Cono, 0.73 ± 0.01, N = 18; SNX, 0.77 ± 0.06, N = 6; NiCl2, 0.78 ± 0.02, N = 9). The combined administration of Aga, Cono, and NiCl2 (3-mix) decreased the mEPSC frequency to 0.47 ± 0.02 (N = 10, Figs. 2Ad-e, C), suggesting that 91% of [Ca^2+^]e-dependent mEPSCs are mediated by VGCCs. In the presence of 3-mix, changing [Ca^2+^]e no longer affected the mEPSC frequency (Figs. 2Ad-e, C), further confirming the role of VGCCs in [Ca^2+^]e-dependent miniature events.

**Fig. 2.**
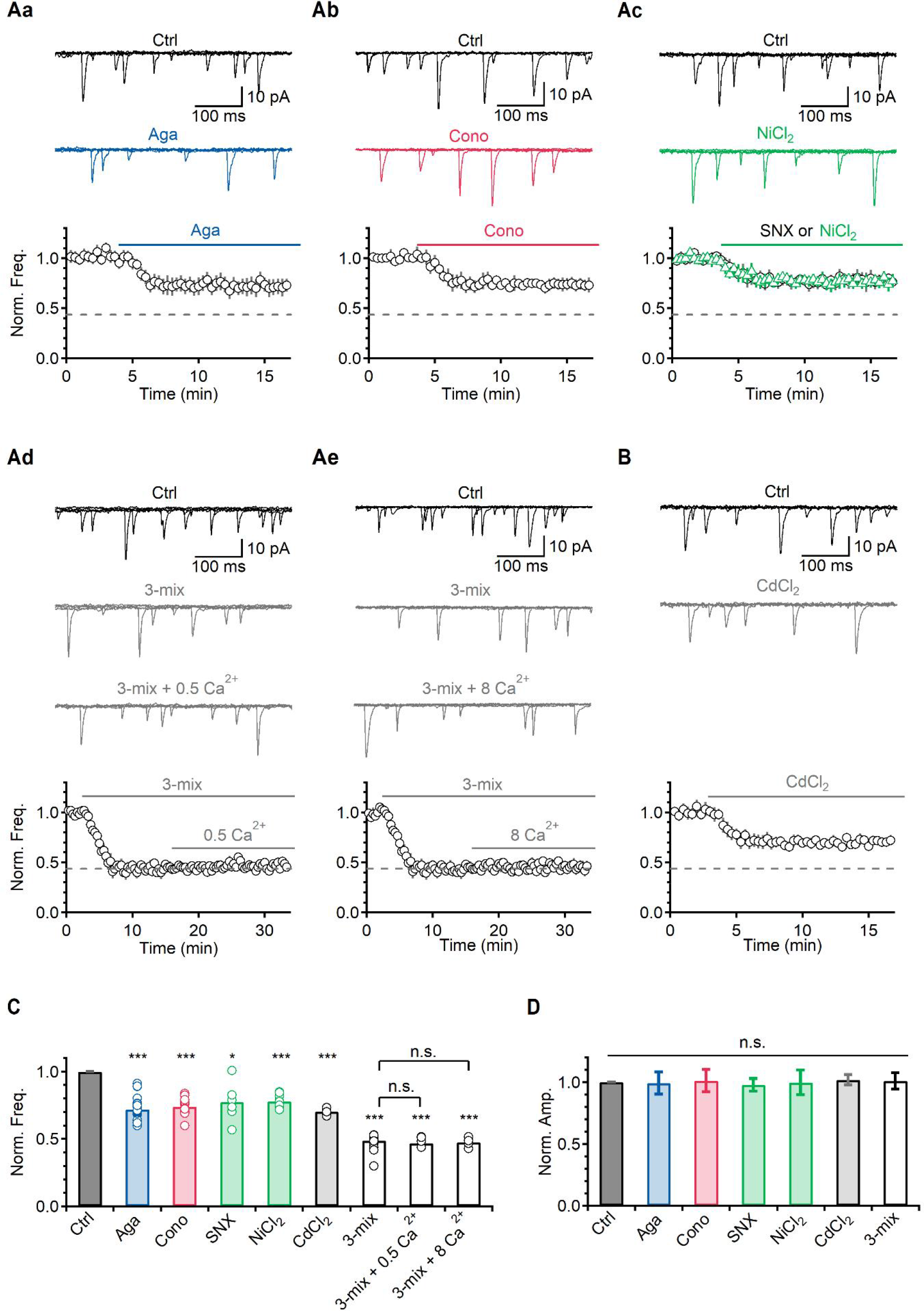
VGCCs contribute to spontaneous glutamatergic release. (A, B) Top. Representative traces of mEPSCs in control and the presence of Aga (Aa), Cono (Ab), NiCl_2_ (Ac), 3-mix (Aga + Cono + NiCl_2_) followed by 0.5 Ca^2+^ (Ad), 3-mix followed by 8 Ca^2+^ (Ae), and CdCl_2_ (B), respectively. Five 500 ms-long mEPSC traces were overlaid. Bottom. Average time courses of the normalized mEPSC frequency. In each time course plot, solid lines indicate the presence of each VGCC blocker. The data were normalized by the mean mEPSC frequency of control. Grey dashed lines (0.42) indicate the division of [Ca^2+^]_e_-dependent and -independent mEPSCs. (C) A bar graph of average values of the normalized mEPSC frequency in different conditions (Aga, 0.71 ± 0.02, *N* = 24, *P*<0.001; Cono, 0.73 ± 0.01, *N* = 18; SNX, 0.77 ± 0.06, *N* = 6; NiCl_2_, 0.78 ± 0.02, *N* = 9; CdCl_2_, 0.70 ± 0.03, *N* = 4; 3-mix, 0.47 ± 0.02, *N* = 10; 3-mix + 0.5 Ca^2+^, 0.47 ± 0.01, *N* = 5; 3-mix + 8 Ca^2+^, 0.48 ± 0.02, *N* = 4; compared to control) (D) A bar graph of average values of the normalized mEPSC amplitude. All data are represented as mean ± S.E.M., \**P*<0.05, \*\*\**P*<0.001, single group mean *t* test or paired *t*-test (C); n.s. = not significant.

One of the strong pieces of evidence supporting the lack of VGCC contribution to spontaneous glutamate release is that Cd^2+^, a nonselective VGCC blocker, has no significant effect (Dai et al., 2015; Tsintsadze et al., 2017; Vyleta and Smith, 2011). We measured mEPSCs in the presence of Cd^2+^ (100 μM) and found that mEPSC frequency was reduced to 0.70 ± 0.03 (N = 4, Figs. 2B, C). This was a significant decrease, yet smaller than that by 3-mix, suggesting that Cd^2+^ did not completely block the VGCC-dependent portion of spontaneous release. This concentration of Cd^2+^ was potent in abolishing depolarization-induced Ca^2+^ currents (Supplementary Fig. 2, 0.04 ± 0.01, N = 3). Since VGCC blockade by Cd^2+^ is voltage-dependent and not as effective at hyperpolarizing potential (Swandulla and Armstrong, 1989), Cd^2+^ may not be suitable for assessing VGCC dependence at the resting membrane potential. We confirmed that mEPSC amplitudes were not changed by any VGCC blocker (Fig. 2D).

To further understand the role of VGCCs in glutamate release, we compared the effects of [Ca^2+^]e and VGCC blockers on eEPSCs with those on mEPSCs. When eEPSCs were recorded at different [Ca^2+^]e, the amplitude of eEPSCs changed according to [Ca^2+^]e, as did the mEPSC frequency (Fig. 3A). The slope of the log-log plot for eEPSC amplitude against [Ca^2+^]e was 1.84 (black line, Fig. 3Ba) which was smaller than that obtained at the neuromuscular junction (3.8) (Dodge and Rahamimoff, 1967), but compatible with the value of 1.77 obtained at hippocampal synapses (Vyleta and Jonas, 2014). Removal of external Ca^2+^ completely abolished eEPSCs (Supplementary Fig. 3A), whereas ryanodine had no effect (Supplementary Fig. 3B), confirming that evoked release was entirely dependent on Ca^2+^ influx via VGCCs. As expected, each VGCC blocker decreased the eEPSC amplitude significantly (Figs. 3C, D), although the contributions of P/Q-, N-, and R-type VGCCs were slightly different from their contributions to mEPSCs (P/Q-type, 65% vs 53%; N-type, 40% vs 49%; R-type, 40% vs 42%; Figs. 2C, 3E). The sum of contributions from the three VGCCs exceeded 100% (145% for eEPSCs and 144% for mEPSCs, Fig. 3E), suggesting that the contribution of each type of VGCC is not entirely independent, but co-activation of multiple VGCCs participates in spontaneous release. The number and organization of VGCCs in the active zone can be evaluated during short-term plasticity in the presence of VGCC blockers. For example, a subtype-specific blockade of VGCCs would not change the paired-pulse ratio (PPR) if vesicle release is triggered by only one type of VGCCs, whereas it would increase the PPR if multiple channel types are involved (Scimemi and Diamond, 2012). Thus, we evaluated changes in the PPR by subtype specific blockers and found that each blocker significantly increased the PPR in association with a decrease of eEPSC amplitude (Fig. 3F). The PPR was unchanged by submaximal concentrations of a single blocker when only a single VGCC subtype remained active in the presence of the other two blockers (Supplementary Fig. 4). These results are compatible with contribution of multiple channel types to vesicular release.

**Fig. 3.**
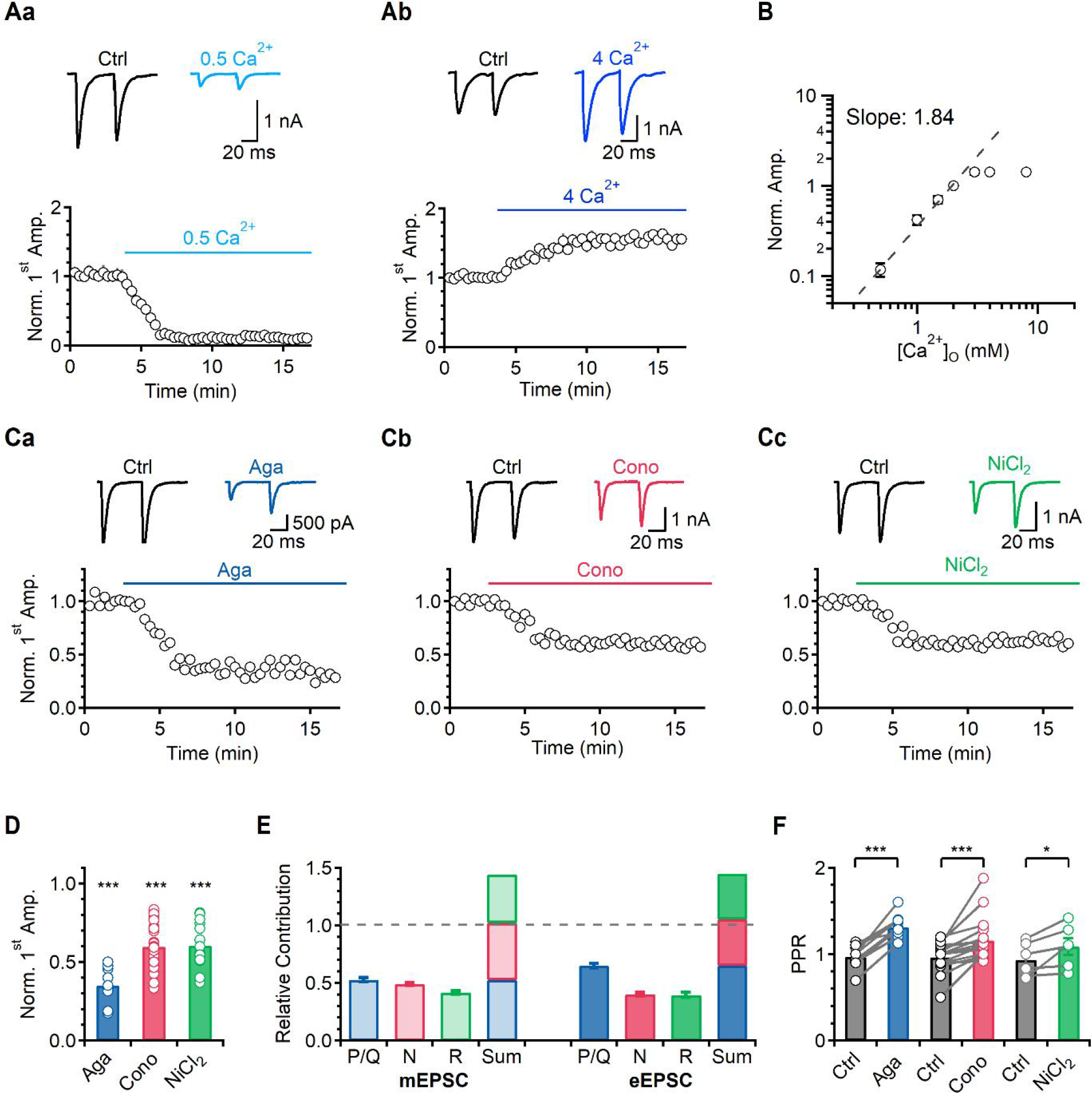
VGCCs contribute to evoked glutamate release. (Aa-b) Top. Representative traces of eEPSC in different [Ca^2+^]_e_ from 2 (Ctrl) to 0.5 mM (0.5 Ca^2+^, light blue) (Aa) and from 2 to 4 mM (4 Ca^2+^, dark blue) (Ab). External [Mg^2+^] was 1 mM for these recordings. Bottom. Average time courses of the normalized 1^st^ eEPSC amplitude (Aa, 0.5 Ca^2+^, 0.11 ± 0.02, *N* = 16; Ab, 4 Ca^2+^, 1.42 ± 0.04, *N* = 6; compared to control, *P*<0.001). In each time course plot, solid lines indicate the presence of 0.5 and 4 Ca^2^, respectively. The data were normalized by the mean amplitude of the 1^st^ eEPSC amplitude of control. (B) A log-log plot for the normalized eEPSC amplitude against [Ca^2+^]_e_. The slope of eEPSC amplitude was 1.84 which was fitted from control (2 mM Ca^2+^) to lower concentration (black circles and dash line). (Ca-c) Top. Representative traces of eEPSCs in control and the presence of Aga (Ca), Cono (Cb), or NiCl_2_ (Cc). Bottom. Average time courses of the normalized 1^st^ eEPSC amplitude. In each time course plot, solid lines indicate the presence of each blocker. The data were normalized by the mean amplitude of the 1^st^ eEPSC amplitude of control. (D) A bar graph of average values of the normalized 1^st^ eEPSC amplitude in different conditions (Aga, 0.35 ± 0.03, *N* = 12; Cono, 0.6 ± 0.02, *N* = 37; NiCl_2_, 0.6 ± 0.03, *N* = 18; compared to control). (E) A chart for the VGCC contribution to mEPSCs (pale bar) or eEPSCs (solid bar). Note that the arithmetic sum of the contributions of each blocker exceeded 1 (grey dashed line) both in spontaneous and evoked release. (F) A bar graph of average values of the paired-pulse ratio (PPR) in different conditions (Ctrl *vs* Aga, 0.97 ± 0.04 *vs* 1.31 ± 0.04; Ctrl *vs* Cono, 0.97 ± 0.04 *vs* 1.16 ± 0.06; Ctrl *vs* NiCl_2_, 0.93 ± 0.04 *vs* 1.09 ± 0.09, paired *t* test). All data are represented as mean ± S.E.M., \**P*<0.05, \*\**P*<0.01, \*\*\**P*<0.001, single group mean *t* test or paired *t*-test (F); n.s. = not significant.

### Nanodomain coupling between VGCCs and Ca^2+^ sensors in autaptic hippocampal neurons

The above results demonstrated that presynaptic VGCCs contribute to both spontaneous and evoked releases in autaptic hippocampal neurons. However, it is not certain whether Ca^2+^ influx upon stochastic openings of VGCCs at resting state directly triggers spontaneous release by increasing local Ca^2+^ levels near primed vesicles at the active zone (Ermolyuk et al., 2013) or indirectly modulates spontaneous release by increasing presynaptic global Ca^2+^ levels. This question can be answered by assessing how tightly vesicular Ca^2+^ sensors for spontaneous release are coupled to Ca^2+^ sources and VGCCs. The distance between VGCCs and Ca^2+^ sensors can be probed using exogenous Ca^2+^ chelators with different kinetics, e.g., EGTA and 1,2-bis(o-aminophenoxy)ethane-N,N,N′,N′-tetraacetic acid (BAPTA) (Adler et al., 1991; Neher, 1998). When the channel-sensor coupling is very tight, the so-called nanodomain coupling, fast Ca^2+^ buffers such as BAPTA, but not slow Ca^2+^ buffers such as EGTA, can interfere with exocytosis, whereas when the coupling is loose, both EGTA and BAPTA can interfere (Adler et al., 1991; Neher, 1998). To estimate the coupling distance of VGCCs with vesicular Ca^2+^ sensors that are involved in spontaneous release, we used pipette solutions containing different concentrations of EGTA and BAPTA. For comparison, we measured mEPSC frequency and eEPSC amplitude from the same cells in such a way that eEPSCs were recorded every 20 s starting immediately after patch break-in, while mEPSCs were recorded continuously between eEPSCs (Fig. 4A). Sequential recordings for eEPSCs and mEPSCs continued until the effects of EGTA or BAPTA perfusion reached a steady state. These experiments were performed in the presence of ryanodine to exclude the effects of exogenous buffers on RyR-dependent mEPSCs (Supplementary Fig. 5). We first confirmed that the eEPSC amplitude in 0.1 mM EGTA remained stable for a recording time of up to 20 min under this experimental protocol (Fig. 4B). The frequency of spontaneous activities increased after eEPSC recording due to asynchronous release, but this increase returned to the control level within a few seconds (Fig. 4Bb). Therefore, we regarded mEPSCs for 10 s before the next stimulation as spontaneous activities. We then examined the effects of increased EGTA or BAPTA concentrations. Changes in eEPSCs or mEPSCs were insignificant with 5 mM EGTA (black symbols, Figs. 4Cb, Db left panel), whereas a significant reduction was induced in both eEPSC amplitude and mEPSC frequency with 5 mM BAPTA (blue symbols, Figs. 4Cb, Db left panel). Differential effects of EGTA and BAPTA suggest a tight coupling between VGCCs and Ca^2+^ sensors for both spontaneous and evoked releases. To estimate the distance between Ca^2+^ sensors and VGCCs, we recorded eEPSCs and mEPSCs with different concentrations of BAPTA and EGTA (Figs. 4Ea-b). Then, we subtracted the [Ca^2+^]e-independent portions (0.26 ± 0.02, N = 5, in 5 mM EGTA and 0.25 ± 0.03, N = 7, in 5 mM BAPTA) from the measured mEPSC frequencies (Db, left panel) to obtain changes in [Ca^2+^]e-dependent mEPSCs (Fig. 4Db, right panel). The resulting curves (orange lines) were superimposed on changes in eEPSC amplitude for comparison (Figs. 4Ea-b). The eEPSC amplitude (green lines) and mEPSC frequency (orange lines) decreased as the concentration of each buffer increased (Figs. 4Ea-b) with time course and magnitude indistinguishable between eEPSCs and mEPSCs at all concentrations tested (Fig. 4E). The distance between Ca^2+^ sources and sensors (r) was estimated by fitting these values to the following equation (see Methods):

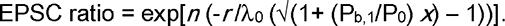

where x is [EGTA] or [BAPTA] in mM; n is the power dependence of vesicle release on [Ca^2+^], λ0 is the length constant of a calcium microdomain in the presence of endogenous buffer (B0) alone; and P0 and Pb,1 are buffer products (kon [B]) of B0 and 1 mM EGTA or BAPTA, respectively. Since decreases in mEPSC frequency and eEPSC amplitude by EGTA and BAPTA were almost identical, we pooled the data for fitting. The best fit was obtained at r = 22 nm with n = 1.62 and P0 = 3000 (Fig. 4F). Taken together, both spontaneous and evoked glutamate release is mediated by nanodomain coupling between VGCCs and vesicular Ca^2+^ sensors in cultured hippocampal neurons.

**Fig. 4.**
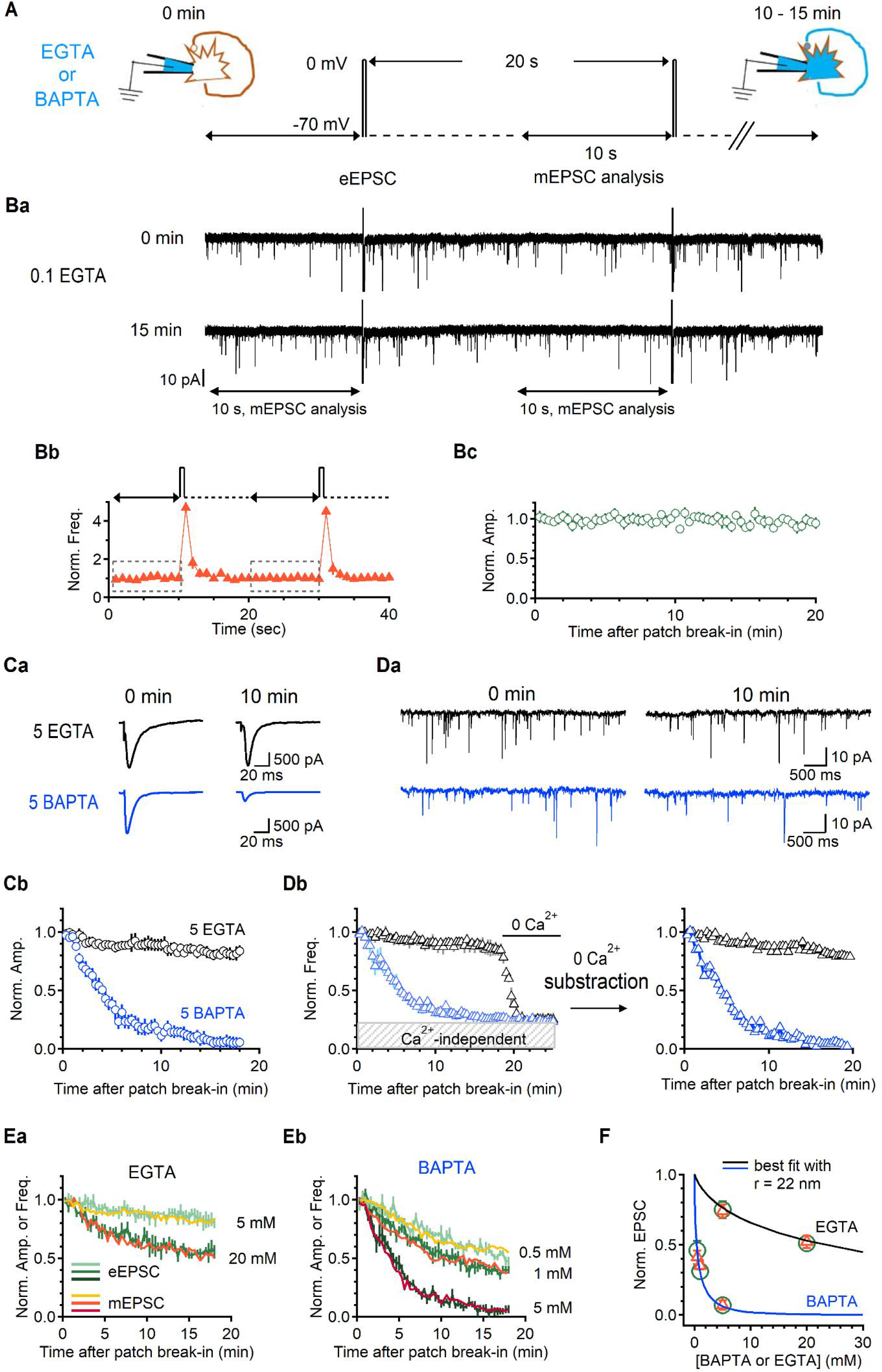
Nanodomain coupling between Ca^2+^ sources and Ca^2+^ sensors for both spontaneous and evoked release in autaptic hippocampal neurons. (A) An experimental scheme for sequential recordings of eEPSCs and mEPSCs from the same cell. (Ba) Representative traces of eEPSCs and mEPSCs at 0 min (top) and 15 min (bottom) after patch break-in with internal 0.1 mM EGTA (0.1 EGTA). (Bb) Top. An experimental scheme for sequential recordings of eEPSCs and mEPSCs. Bottom. An average time course of the mEPSC frequency normalized by the data acquired from initial 10s with 1 s bin. Dashed-boxes indicate the mEPSC frequency analysis sections. (Bc) An average sequential time plot of the eEPSC amplitude normalized by the data acquired from initial 60s. (Ca) Representative traces of eEPSCs at 0 min and 10 min after patch break-in with internal 5 mM EGTA (5 EGTA; black) or 5 mM BAPTA (5 BAPTA; blue). (Cb) Average time courses of the normalized eEPSC amplitude in 5 EGTA and 5 BAPTA. The data were normalized by the mean amplitude of the first 3 eEPSCs after patch break-in. The magnitude of the reduction at steady state and the time course (τ) were 0.75 ± 0.06 *vs* 0.07 ± 0.05 and 1146 ± 85 s *vs* 221 ± 9.2 s (5 EGTA *vs* 5 BAPTA; *N* = 5 *vs* 7). (Da) Representative traces of mEPSCs at 0 min and 10 min after patch break-in in 5 EGTA and 5 BAPTA. (Db) Left. Average time courses of the normalized mEPSC frequency. The data were normalized by the mean frequency of the initial 1 min after patch break-in. When the response reached to the steady state, extracellular solution was changed to 0 Ca^2+^ to measure Ca^2+^-independent mEPSCs. Right. [Ca^2+^]_e_-dependent mEPSC frequency was calculated by subtracting the Ca^2+^-independent portion from the total mEPSCs. The magnitude of the reduction at steady state and the time course (τ) were 0.74 ± 0.05 *vs* 0.07 ± 0.03 and 1004 ± 107 s *vs* 277 ± 12.3 s (5 EGTA *vs* 5 BAPTA). (Ea-b) Average sequential time plots for the normalized mEPSC frequency (light orange, orange, and dark orange) or the normalized eEPSC amplitude (light green, green, and dark green) in the presence of varying concentrations of Ca^2+^ chelators in the intracellular solution (Ea, 20 EGTA, eEPSC *vs* mEPSC, 0.51 ± 0.03 *vs* 0.52 ± 0.04 and 307.7 ± 10.1 s *vs* 363.6 ± 13.4 s, steady state and τ, *N* = 4; Eb, 0.5 BAPTA, 0.46 ± 0.07 *vs* 0.41 ± 0.05 and 588.2 ± 37.6 s *vs* 625 ± 40 s, steady state and τ, *N* = 5; 1 BAPTA, 0.31 ± 0.04 *vs* 0.35 ± 0.03 and 357.1 ± 11.8 s *vs* 335.8 ± 14.3 s, steady state and τ, *N* = 8). (F) Concentration-effect curves for EGTA (black) and BAPTA (blue). Symbols indicate the steady state values obtained in Ea and Eb (orange triangles for mEPSCs; green circles for eEPSCs). Lines are best fit curves for the steady state value data obtained with EGTA and BAPTA using the equation, EPSC ratio = exp[ n (-r /γ_0_ (√(1+ (P_b,1_/P_0_) x) - 1))], described in Methods. All data are represented as mean ± S.E.M.

### The coupling distance between VGCCs and Ca^2+^ sensors determines the contribution of VGCCs to spontaneous release at calyx of Held synapses

Previous studies reported that only spontaneous GABA release, but not spontaneous glutamate release, is mediated by VGCCs (Courtney et al., 2018; Tsintsadze et al., 2017; Vyleta and Smith, 2011), whereas Ermolyuk et al. (2013) and our results showed that VGCCs contribute to spontaneous glutamate release in hippocampal neuron cultures. These results imply that VGCCs contribute to glutamate release only under certain circumstances. We hypothesized that stochastic opening of VGCCs can trigger exocytosis only when VGCCs and vesicular Ca^2+^ sensors are very close to nanodomain coupling. To test this hypothesis, we investigated the contribution of VGCCs to mEPSCs at calyx of Held synapses, where the coupling distance between VGCCs and vesicular Ca^2+^ sensors changes from loose (∼100 nm) to tight coupling (∼20 nm) during development (Fedchyshyn and Wang, 2005; Wang et al., 2009). The mEPSC frequency at immature synapses was [Ca^2+^]e-dependent, yet not mediated by VGCCs (3-mix, 0.94 ± 0.09; 0 mM [Ca^2+^]e, 0.43 ± 0.04, N = 13, Figs. 5Aa-b), which is consistent with previous reports (Dai et al., 2015; Tsintsadze et al., 2017). All recordings were performed within 10 min after break-in to avoid any rundown. Recent studies have shown that the mature calyx of Held is morphologically heterogeneous and is grouped into three types depending on the number of swellings (type 1, less than 5 swellings; type 3, more than 15 swellings; type 2, in between; Grande and Wang, 2011). In addition, type 1 calyces contain synaptic vesicles tightly coupled to VGCCs (∼14 nm, stalk), whereas type 3 calyces harbor a mixture of loosely (30-50 nm, swellings) and tightly coupled vesicles (Fekete et al., 2019). Because VGCC contribution to mEPSCs may vary depending on calyx maturity, we analyzed 3-mix effects on mEPSC frequency for each type separately. For this, after recordings of mEPSCs from postsynaptic neurons of the medial nucleus of the trapezoid body, pairing presynaptic calyx terminals were filled with Alexa Fluor 488 using another whole-cell pipette. High-resolution confocal Z-stack images (used for 3D reconstruction of the calyx structure) of the recorded calyces were obtained after careful withdrawal of the presynaptic pipette to allow resealing of the presynaptic membrane. All mEPSC frequencies at mature synapses were [Ca^2+^]e-dependent and showed significant inhibition by 3-mix, but inhibition was greatest for type 1 (3-mix, 0.32 ± 0.02, 0.62 ± 0.03, and 0.79 ± 0.02; 0 mM [Ca^2+^]e, 0.16 ± 0.01, 0.17 ± 0.01, and 0.12 ± 0.01; N = 13, 12, and 11 for type 1, 2, and 3, respectively; Figs. 5Bb to Db). Mature calyx synapses showed a higher basal mEPSC frequency (∼2 Hz) compared to immature calyx synapses (∼0.5 Hz). Despite their different sensitivities to 3-mix, the basal mEPSC frequency did not differ among morphologically distinct types (Figs. 5Ea-b). The mEPSC frequency was similar for all three types under 0 mM [Ca^2+^]e and lower than that of immature calyx synapses, suggesting that the [Ca^2+^]e- dependent portion of mEPSCs was greater for mature synapses (Fig. 5Eb). The [Ca^2+^]e- and VGCC-dependent relative contributions to mEPSC frequency for all recorded calyx synapses are summarized in Figs. 5Ec, Ed. Considering that P/Q-type calcium channels predominate in mature calyx terminals (Iwasaki and Takahashi, 1998), our data suggest that VGCCs are engaged in spontaneous release during synaptic maturation by tightening the coupling between VGCCs and Ca^2+^ sensors, especially in mature type 1 calyx synapses. This is similar to the VGCC-dependent spontaneous GABA release in previous studies (Courtney et al., 2018; Tsintsadze et al., 2017; Vyleta and Smith, 2011) and thus suggests that tight coupling of VGCCs and vesicular Ca^2+^ sensors is a prerequisite for VGCC-dependent spontaneous release.

**Fig. 5.**
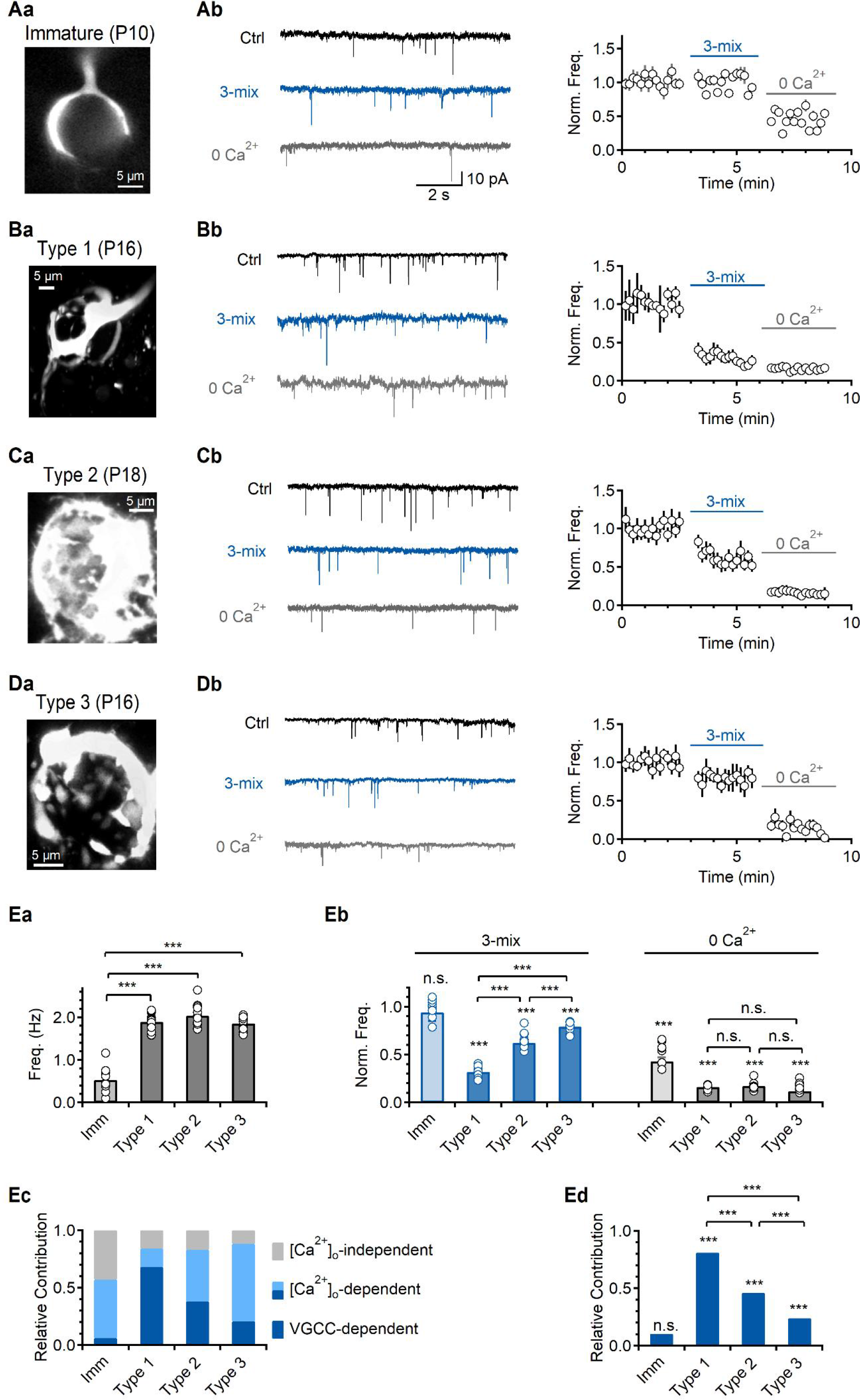
Contribution of VGCCs to spontaneous release at the calyx of Held synapses. (A-D) The calyx of Held synapses at different developmental stages. (Aa-Da) 3D reconstruction images of immature (P10, Aa), type 1 (P16, Ba), type 2 (P18, Ca), or type 3 (P16, Da) calyces filled with Alexa-488. (Ab-Db) Left. Representative traces of mEPSC in control and the presence 3-mix (blue) and 0 Ca^2+^ (grey) at immature (Ab; P7 - 10) or mature (Bb-Db; P16 - 18) calyx of Held synapses. Right. Average time courses of the normalized mEPSC frequency. In each time course plot, solid lines indicate the presence of 3-mix or 0 Ca^2+^. The data were normalized by the mean frequency of control. (Ea) A bar graph of average values of mEPSC frequency at immature and mature synapses (immature, 0.53 ± 0.09 Hz, *N* = 13; type 1, 1.89 ± 0.05 Hz, *N* = 13; type 2, 2.03 ± 0.06 Hz, *N* = 12; type 3, 1.86 ± 0.04 Hz, *N* = 11). (Eb) A bar graph of average values of the normalized mEPSC frequency in different conditions (3-mix, immature, 0.94 ± 0.09; type 1, 0.32 ± 0.02; type 2, 0.62 ± 0.03; type 3, 0.79 ± 0.02; 0 Ca^2+^, immature, 0.43 ± 0.04; type 1, 0.16 ± 0.01; type 2, 0.17 ± 0.01; type 3, 0.12 ± 0.01). (Ec) A summary bar graph of the relative contribution to Ca^2+^- dependence on mEPSCs (immature, 0.57; type 1, 0.84; type 2, 0.83; type 3, 0.88). (Ed) A summary bar graph of average values of VGCC contribution to Ca^2+^-dependence mEPSCs (immature, 0.1; type 1, 0.81; type 2, 0.46; type 3, 0.24). All data are represented as mean ± S.E.M., \**P*<0.05, \*\**P*<0.01, \*\*\**P*<0.001, single group mean *t*-test or paired *t*-test (E); n.s. = not significant.

### Calmodulin mediates VGCC-dependent spontaneous release at mossy fiber CA3 synapses

We then asked whether the contribution of VGCCs to spontaneous release also applies to different glutamatergic synapses by examining the effect of VGCC blockers on mEPSCs in CA1-PCs and CA3-PCs. The basal mEPSC frequency was higher in CA3 (2.2 ± 0.30, N = 24) than in CA1 (1.01 ± 0.07, N = 15), yet the [Ca^2+^]e-dependent portion was not substantially different (0.54 ± 0.04 vs 0.58 ± 0.03, Figs. 6Aa-c). However, administration of 3-mix significantly decreased the mEPSC frequency in CA3-PCs, but not in CA1-PCs (Figs. 6Aa-c), indicating that the contribution of VGCCs to spontaneous release is different between the two types of neurons. To examine whether the difference in coupling distance between VGCCs and vesicular sensors underlies the difference in VGCC dependence of spontaneous release, we compared the effects of EGTA-AM on eEPSCs evoked by stimulating major inputs to each neuron, i.e., Shaffer collaterals (SC) to CA1-PCs and mossy fibers (MF) to CA3-PCs. VGCC dependence of mEPSCs in CA3-PCs. We, therefore, hypothesized the presence of other mechanisms besides coupling distance whereby calcium entry through VGCCs located remotely from calcium sensors can contribute to exocytosis. As a possible mechanism, we examined the role of calmodulin, a Ca^2+^-binding protein that can modulate neurotransmission, in regulating VGCC-dependence of spontaneous release. The application of 10 μM calmidazolium, a calmodulin inhibitor, significantly reduced mEPSC frequency in CA3-PCs, but not in CA1-PCs (Figs 6Ca-d), suggesting that the role of calmodulin in spontaneous release is specific to CA3-PCs. In the presence of calmidazolium, no further reduction was observed in mEPSC frequency by subsequent application of 3-mix (Fig. 6Cb), indicating that calmidazolium inhibited VGCC-dependent spontaneous release. These results suggest that calmodulin mediates VGCC-dependent spontaneous release at loosely coupled CA3 synapses.

**Fig. 6.**
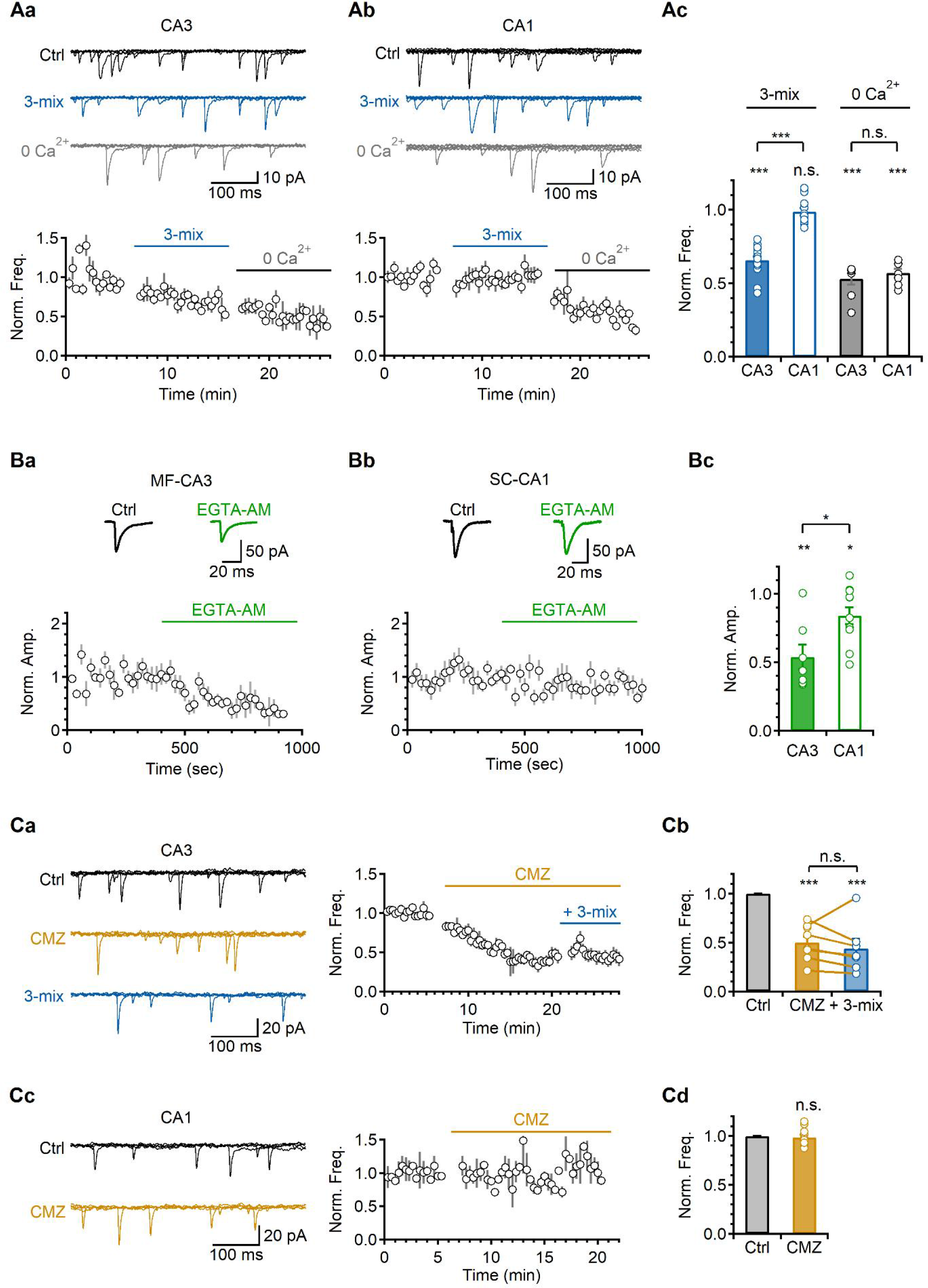
Calmodulin mediates VGCC-dependent spontaneous release in CA3-PCs. (Aa-b) Top. Representative traces of mEPSCs in control and the presence of 3-mix and 0 Ca^2+^ in CA3 (Aa) and CA1 (Ab). Five 500 ms-long mEPSC traces were overlaid. Bottom. Average time courses of the normalized mEPSC frequency. In each time course plot, solid lines indicate the presence of 3-mix or 0 Ca^2+^. The data were normalized by the mean frequency of control. (Ac) A bar graph of average values of the normalized mEPSC frequency in different conditions (3-mix, CA3, 0.65 ± 0.02, *N* = 16; CA1, 0.99 ± 0.02, *N* = 11; 0 Ca^2+^, CA3, 0.53 ± 0.04, *N* = 8; CA1, 0.57 ± 0.03, *N* = 9). (Ba-b) Top. Representative traces of eEPSCs in control and the presence of 50 μM EGTA-AM (green) in CA3 (Ba) and CA1-PCs (Bb) in response to MF and SC stimulation, respectively. Bottom. Average time courses of the normalized eEPSC amplitude. In each time course plot, solid lines indicate the presence of 50 μM EGTA-AM. The data were normalized by the mean eEPSC amplitude of control. (Bc) A bar graph of average values of the normalized eEPSC amplitude in different condition (CA3, 0.54 ± 0.09, *N* = 7, *P* = 0.003; CA1, 0.84 ± 0.06, *N* = 11, *P* = 0.028). (Ca) Left. Representative traces of mEPSCs in control and the presence of CMZ (yellow) and CMZ with 3-mix (blue) in CA3. Right. Average time courses of the normalized mEPSC frequency. Solid lines indicate the presence of drugs. (Cb) A bar graph of average values of the normalized mEPSC frequency in different conditions (CMZ, 0.50 ± 0.03, *N* = 9; CMZ + 3- mix, 0.44 ± 0.06, *N* = 7). (Cc) Left. Representative traces of mEPSCs in control and the presence of CMZ (yellow) in CA1. Right. Average time courses of the normalized mEPSC frequency. Solid line indicates the presence of CMZ. (Cd) A bar graph of average values of the normalized mEPSC frequency in CMZ compared to control (0.99 ± 0.04, *N* = 4). All data are represented as mean ± S.E.M., \**P*<0.05, \*\**P*<0.01, \*\*\**P*<0.001, single group mean *t*-test or paired *t*-test (Bc, Cb); n.s. = not significant.

## DISCUSSION

While Ca^2+^ influx through VGCCs in presynaptic axon terminals is critical for evoked synaptic transmitter release, the role of VGCCs in spontaneous release at glutamatergic synapses remains controversial (Williams and Smith, 2018). We investigated this issue in various glutamatergic synapses and obtained distinct results among different synapses. We found that mEPSCs were indeed dependent on Ca^2+^ influx via P/Q-, N-, and R-type VGCCs in cultured autaptic hippocampal neurons and mature calyx of Held where coupling between VGCCs and vesicular Ca^2+^ sensors was estimated (22 nm in autaptic neurons; Fig. 4F) or reported to be very tight (14 nm in type I calyx of Held; Fekete et al., 2019), On the other hand, the VGCC dependence of spontaneous release is variable at loosely coupled synapses. VGCC-dependent spontaneous release does not exist in the immature calyx of Held and CA1-PCs but occurs via calmodulin in CA3-PCs. Taken together, our data strongly suggest that the coupling distance between VGCCs and Ca^2+^ sensors is a critical determinant in the VGCC dependence of spontaneous release, yet other Ca^2+^-dependent signaling mechanisms such as calmoduin can be involved in specific synapses. Our data suggest distinct roles and regulatory mechanisms of spontaneous release in various types of synapses.

It has been of great interest whether the molecular machineries known for evoked release, such as Ca^2+^ sources, Ca^2+^ sensors, vesicle pools, also operate for Ca^2+^-dependent spontaneous release (Kaeser and Regehr, 2014; Schneggenburger and Rosenmund, 2015). Ca^2+^ cooperativity of mEPSCs was reported to be lower than that of eEPSCs (Vyleta and Smith, 2011), and this low Ca^2+^ sensitivity was regarded to represent the involvement of different mechanisms between spontaneous and evoked release. In the present study, we used cultured autaptic neurons where mEPSCs and eEPSCs are originated from the same set of presynaptic terminals, and thus, direct comparison of the two are allowed. We obtained Ca^2+^-sensitivity for mEPSCs from the [Ca^2+^]e-dependent portion of mEPSCs, calculated from observed mEPSCs by subtracting [Ca^2+^]e-independent mEPSCs, and found that it was comparable to Ca^2+^ sensitivity of eEPSCs (Figs 1, 3). The sensitivity to EGTA or BAPTA for [Ca^2+^]e-dependent mEPSCs was also not significantly different from that for eEPSCs (Fig. 4). Moreover, [Ca^2+^]e-dependent spontaneous release was mediated by P/Q-, N-, and R-type VGCCs as evoked release (Figs. 2, 3). These results suggest that the underlying mechanisms of [Ca^2+^]e-dependent spontaneous release may not be substantially different from those of evoked release in cultured autaptic neurons. However, we also found that spontaneous release in CA1-PCs and immature calyx of Held is not VGCC-dependent (Figs. 5, 6), suggesting that [Ca^2+^]e-dependent mechanisms for spontaneous glutamate release are not universal among different types of synapses. The question of whether spontaneous and evoked release originate from the same or separate pools of vesicles is also controversial (Chung et al., 2010; Fredj and Burrone, 2009; Groemer and Klingauf, 2007; Hua et al., 2010). Diverse views about the molecular mechanisms of spontaneous release may suggest that spontaneous release is not governed by a single mechanism, but is attributable to multiple pathways.

Contribution of VGCCs to spontaneous release was well characterized at GABAergic synapses in cortical cultures (Williams et al., 2012; Williams and Smith, 2018), but it was disputed in glutamatergic synapses in cultured cortical neurons (Courtney et al., 2018; Vyleta and Smith, 2011), CA1 hippocampal neurons (Babiec and O’Dell, 2018), and immature calyx of Held (Dai et al., 2015; Tsintsadze et al., 2017). Therefore, a lack of VGCC contribution was regarded to represent fundamental differences in molecular mechanisms between GABAergic and glutamatergic synapses (Williams and Smith, 2018). However, we showed that VGCC contribution is variable among different glutamatergic synapses, revealing that the lack of VGCC contribution to spontaneous release is not a general feature of glutamatergic synapses. We showed VGCC contribution in hippocampal cultures and calyx of Held that have tightly coupled synapses (Figs. 2, 4, 5), while the lack of VGCC contribution to spontaneous release in immature calyx of Held and CA1-PC synapses (Figs. 5, 6), which are known as loosely coupled synapses (Blatow et al., 2003). More general notion may be that VGCCs contribute to spontaneous release at tightly coupled synapses in a way that stochastic opening of VGCCs directly trigger exocytosis.

In spite that we and others reported a lack of VGCC contribution to spontaneous release in most of excitatory neurons with loosely coupled synapse, CA3-PCs revealed an exception. VGCC contribution to spontaneous release at loosely coupled synapses was also reported at GABAergic synapses in dentate granule cells (Goswami et al., 2012). We found that the mEPSC frequency was reduced by calmidazolium in CA3-PCs, but not in CA1-PCs (Fig. 6), suggesting that calmodulin plays an active role in regulating spontaneous release at inputs to CA3-PCs, but not to CA1-PCs. Since VGCC blockers did not affect mEPSCs in CA3-PCs in the presence of calmidazolium, we concluded that the VGCC dependence of spontaneous release in CA3-PCs is mediated by calmodulin. Calmodulin regulates calcium channel activity and plays a role in synaptic vesicle release and recycling(Ben-Johny and Yue, 2014; Liang et al., 2021; Sakaba and Neher, 2001). The role of calmodulin in spontaneous release has not been reported in mammalian central synapses; however, calmodulin participates in spontaneous release at various synapses such as the Drosophila neuromuscular junction and rat retinal ribbon synapses through the v-ATPase subunit and myosin light chain kinase, respectively (Liang et al., 2021; Wang et al., 2014). The involvement of calmodulin implies that spontaneous release is not a passive phenomenon determined by the physical location of VGCCs and synaptic vesicles but can be modulated by physiological conditions that change the activity of calmodulin. However, we do not know as to why calmodulin plays a role in spontaneous release specifically in CA3-PCs or what the mechanism and physiological significance of calmodulin-dependent regulation of spontaneous release is. These questions remain to be investigated in future studies. It also needs to be elucidated whether calmodulin is also involved in the contribution of VGCCs to spontaneous release at loosely coupled GABAergic synapses.

The physical distance between presynaptic VGCCs and Ca^2+^ sensors, which may vary depending on the cell types and developmental states (Baur et al., 2015; Bornschein et al., 2019; Taschenberger et al., 2002; Wang et al., 2008), is a key factor for determining the characteristics of AP-triggered neurotransmission (Eggermann et al., 2012; Rebola et al., 2019). A modeling study has revealed that nanodomain coupling offers several functional advantages, including increased synaptic efficacy and speed of synaptic transmission (Bucurenciu et al., 2008). Our study highlights the role of the coupling distance in the regulation of spontaneous neurotransmission. In fact, spontaneous release by stochastic Ca^2+^ channel opening in synapses with nanodomain coupling was considered a disadvantage because it could lead to excessive spontaneous release (Eggermann et al., 2012; Stanley, 1997). However, there was no evidence supporting the occurrence of spontaneous release at higher frequencies in nanodomain synapses compared to microdomain synapses. The mEPSC frequency in CA3-PCs was similar to that in mature calyx of Held synapses (2.2 vs 1.9 Hz, respectively). The mEPSC frequency varied widely between 0.5 and 10 Hz in cultured autaptic neurons (Fig. 1Ab), but the coupling distance was estimated to be invariably short (Fig. 4). These results suggest that there may be a mechanism to protect synapses from excessive spontaneous release generated by stochastic VGCC opening in nanodomain synapses. It has been suggested that the use of two or three open channels rather than a single channel for vesicle release may underlie this protective role (Bucurenciu et al., 2010). We found that both spontaneous and evoked releases were mediated by the co-activation of multiple types of VGCCs in autaptic hippocampal cultures (Figs. 2, 3), which could support this prediction.

Regarding vesicular Ca^2+^ sensors for spontaneous release, the difference between glutamatergic and GABAergic synapses was reported. At inhibitory neurons in cortical neuron culture where the VGCC dependence of spontaneous release is well-characterized (Goswami et al., 2012; Williams et al., 2012), the low-affinity Ca^2+^ sensor synaptotagmin 1 (Syt1) mediated spontaneous release as well as evoked release (Courtney et al., 2018; Xu et al., 2009). VGCC blockers affected Syt1-dependent spontaneous release, but, not in glutamatergic synapses where mEPSCs were independent of VGCCs (Courtney et al., 2018). These findings were regarded to represent divergence in the release machinery between GABAergic and glutamatergic neurons. However, Syt1 contribution to spontaneous release may not be exclusive for GABAergic synapses, but for tightly coupled synapses where spontaneous release is mediated by local Ca^2+^ increases via VGCC activation. To clarify this issue, it needs to be investigated whether Syt1 contributes to spontaneous glutamate release in hippocampal neuron cultures where spontaneous release is VGCC-dependent.

Molecular mechanisms underlying VGCC-independent spontaneous release are less understood. VGCC-independent spontaneous release was shown to be mediated by Doc2, a high-affinity Ca^2+^ sensor (Groffen et al., 2010), where Doc2α and Doc2β contribute to excitatory and inhibitory transmissions, respectively (Courtney et al., 2018). The Ca^2+^ source for Doc2 activation is not clear. Possibly, multiple Ca^2+^ sources involved in the regulation of resting Ca^2+^ levels, such as internal Ca^2+^ sources (Carter et al., 2002; Emptage et al., 2001; Sharma and Vijayaraghavan, 2003) or Ca^2+^ influx via transient receptor potential cation channels (TRPCs) (Peters et al., 2010; Shoudai et al., 2010), may contribute to VGCC-independent spontaneous release by activating Doc2 (Groffen et al., 2010); however, the relationship between these sources and Doc2 remains to be uncovered. Ca^2+^ sensing receptor (CaSR), a G protein-coupled receptor activated by extracellular Ca^2+^, was proposed to mediate VGCC-independent spontaneous release in glutamatergic neurons (Vyleta and Smith, 2011), but how CaSR activation leads to vesicle exocytosis remains elusive.

Taken together, with extensive analysis about the role of VGCCs in spontaneous glutamate release, our study revealed complex natures of spontaneous release that involve distinct Ca^2+^ sources and different regulating mechanisms depending on cell type and condition. It remains to be investigated whether VGCC-dependence has any implication about physiological roles of spontaneous release. Since VGCC-dependent spontaneous release is likely to share common mechanism with evoked release, any changes in presynaptic terminals that affect evoked release could regulate VGCC-dependent spontaneous release, and may not influence VGCC-independent spontaneous release.

## Materials and Methods

### Autaptic Neuronal culture

All preparations were carried out under the animal welfare guideline of Seoul National University (SNU), and approved by IACUC of SNU. Primary cultures of rat hippocampal neurons were prepared as described previously with slight adaptations (Bekkers and Stevens, 1991). Briefly, hippocampal neurons and astrocytes were obtained from Sprague-Dawley (SD) rats according to the protocols approved by the Seoul National University Institutional Animal Care and Use Committee. Astrocyte cultures were prepared from the Sprague-Dawley rat cortices P0 - P1 and grown for 10 days in 100- mm culture dish in glial medium [minimum essential medium (MEM; Invitrogen) supplemented with 0.6 % glucose, 1 mM pyruvate, 2 mM GlutaMAX-I (Invitrogen), 10 % horse serum (HS; Invitrogen), and 1 % penicillin-streptomycin (PS; Invitrogen)] before plating on the sprayed microisland coverslips in 30-mm petri dishes. 2 - 3 days before neurons being added in sprayed microisland dishes, astrocytes were removed from the 100-mm culture dish using trypsin-EDTA (Invitrogen) and plated on the microisland coverslips at a density of 60,000 cells/dish. Hippocampi from P0 - P1 SD rats were dissected in Hank’s balanced salt solution (Invitrogen), digested with papain (Worthington, Freehold, NJ, USA), and then triturated with a polished half-bore Pasteur pipette. Immediately after removing glia medium in 30-mm dishes of microisland-shaped astrocytes, hippocampal neurons were added at a density of 6,000 cells/dish and were grown in neurobasal medium supplemented with B27 and glutamax (Invitrogen).

### Hippocampal Slice Preparation

Hippocampal slices were prepared from 3-week-old (P17 - 21) or 4-week-old (P23 - 30) Sprague-Dawley rats. After anaesthetized by inhalation with 5% isoflurane, rats were decapitated and the brain was quickly removed and chilled in an ice-cold high-magnesium cutting solution containing the following (in mM): 110 choline chloride, 25 NaHCO3, 20 glucose, 2.5 KCl, 1.25 NaH2PO4, 1 sodium pyruvate, 0.5 CaCl2, 7 MgCl2, 0.57 ascorbate (pH 7.3 bubbled with 95 % O2 - 5 % CO2; osmolarity ∼300 mOsm. The isolated brain was glued onto the stage of a vibrating blade microtome (VT1200S, Leica Microsystems) and 300 μm-thick transverse hippocampal slices were cut. The slices were incubated at 34°C for 30 min in artificial cerebrospinal fluid (aCSF) containing the following (in mM): 125 NaCl, 25 NaHCO3, 20 glucose, 2.5 KCl, 1.25 NaH2PO4, 1 sodium pyruvate, 2 CaCl2, 1 MgCl2, 0.57 ascorbate, bubbled with 95 % O2 - 5 % CO2, and thereafter maintained at room temperature until required.

### Calyx of Held Brain Slice Preparation

Transverse brainstem slices (300 μm thick) were prepared from 7 to 18-day-old (P7 - 9, immature pre-hearing onset; P16 - 18, mature post-hearing onset) Sprague-Dawley rats. The standard extracellular solution contained (in mM) 125 NaCl, 2.5 KCl, 25 glucose, 25 NaHCO3, 1.25 NaH2PO4, 0.4 ascorbic acid, 3 myoinositol, 2 Na-pyruvate and 2 CaCl2, 1 MgCl2 (pH 7.4, bubbled with 95 % O2 - 5 % CO2; osmolarity ∼320 mOsm). For slicing, CaCl2 and MgCl2 were changed to 0.5 and 2.5 mM, respectively. Slices were incubated at 34°C for 30 min in the extracellular solution and thereafter maintained at room temperature until required.

### Electrophysiology

For recordings in cultured autaptic neurons, cells were visualized with an Olympus IX70 inverted microscope. Whole-cell voltage- or current-clamp recordings from hippocampal autaptic pyramidal neurons were performed at room temperature and continuously perfused with extracellular solution consisting of following composition (in mM): 135 NaCl, 2.5 KCl, 2 CaCl2, 1 MgCl2, 10 glucose, 10 HEPES, 0.1 EGTA, pH 7.3 adjusted with NaOH (295 - 300 mOsm), and maintained at 0.5 - 1 ml min^-1^. All recordings were done at least 23 days after neurons were plated on coverslips. The internal pipette solution for recording mEPSCs and resting membrane potential (RMP) was used K-gluconate based solution. The unit of mEPSC frequency is Hertz.

For recordings in hippocampal slices, slices were transferred to an immersed recording chamber continuously perfused with oxygenated aCSF using a peristaltic pump (Gilson). CA3 or CA1 pyramidal cells were visualized using an upright microscope equipped with differential interference contrast optics (BX51WI, Olympus). Whole-cell voltage- or current-clamp recordings were performed at 32 ± 1°C and the rate of aCSF perfusion was maintained at 1 - 1.5 ml min^-1^. For high [K^+^]o experiments, NaCl was reduced to maintain osmolarity. Recordings were made in somata with an EPC-10 amplifier (HEKA Electonik, Lambrecht/Pfalz, Germany). Signals were low-pass filtered at 5 kHz (low-pass Bessel filter) and sampled at 10 kHz. Series resistance (Rs) was monitored, and only recordings with Rs remained constant (<30% change during a recording) were used. Rs was compensated to 50 - 70 %. The data were analyzed using IGOR software (Wavemetrics, Lake Oswego, OR, USA). Patch electrodes were pulled from borosilicate glass capillaries to a resistance between 3 and 4 MΩ when filled with pipette solution. The internal pipette solution for recording miniature excitatory postsynaptic currents (mEPSCs) contained the following composition (in mM): 130 Cs-methanesulfate, 8 NaCl, 4 MgATP, 0.3 Na2GTP, 10 HEPES, 0.1 EGTA, pH 7.3 adjusted with CsOH (295 - 300 mOsm). For RMP recording, K-gluconate was substituted for Cs-methanesulfate, and pH was adjusted with KOH.

The mEPSCs of hippocampal slices or hippocampal autapses were recorded at holding potential of - 70 mV. 0.5 μM TTX and 0.1 mM picrotoxin was added during recordings in acute slices. Events exceeding 6 - 7 pA within a specified interval of three to four digitized points (0.5 - 0.8 ms) that showed a single exponential decay time course were identified as mEPSC. The rise time of mEPSC indicated the 20 - 80 % rise time. mEPSC frequency was measured within 20 s bins. Synaptic activities were recorded at a holding potential of -70 mV. eEPSCs were recorded every 20 s after applying depolarization pulses from -70 to 0 mV for 2 ms. From the continuous recordings at -70 mV without stimulations, toxins and chemicals were typically applied for 5 - 20 min until a constant effect was observed. To rule out any effect of Ca^2+^ depletion of internal Ca^2+^ stores on spontaneous release, recordings were completed within 10 min after break-in.

For recording from calyx of Held synapses, brainstem slices were transferred to a recording chamber in an upright microscope and whole-cell patch-clamp recordings were made at room temperature using an EPC10/2 amplifier (HEKA, Pfalz, Germany). Postsynaptic cells were whole-cell voltage clamped at a holding voltage of -70 mV using a pipette (3.0 - 3.5 MΩ) solution containing the following (in mM): 140 Cs-gluconate, 20 tetraethylammonium (TEA)-Cl, 5 Na2-phosphocreatine, 4 MgATP, 0.3 Na2GTP, 20 HEPES, 10 EGTA at pH 7.4 (adjusted with CsOH). Experiments were discarded when series resistance exceeded 10 MΩ (range, 4 - 10 MΩ). 1 μM tetrodotoxin, strychnine, and 100 μM picrotoxin were added during recordings in acute slices. After recordings of mEPSCs from postsynaptic MNTB neurons were complete, pairing presynaptic calyces terminals were filled with Alexa Fluor 488 for less than 1 min using another whole-cell pipette, and then the presynaptic pipette was carefully withdrawn to allow resealing of the presynaptic membrane. We immediately took high-resolution confocal Z-stack images of the recorded calyces, from which the 3D structure of the calyx was reconstructed using FIJI (Grande and Wang, 2011). For this procedure, presynaptic patch electrodes with resistance of 3 to 4 MΩ contained the following (in mM): 140 K-gluconate, 20 KCl, 10 HEPES, 5 Na2-phosphocreatine, 4 Mg-ATP, 0.3 Na-GTP, 0.5 EGTA, and 0.25 Alexa Fluor 488 hydrazide (Invitrogen) with pH adjusted at 7.4.

### Drugs

ω-Agatoxin-IVA, ω-Conotoxin GVIA, SNX-482, TTX, and ryanodine were purchased from Alomone Labs (Jerusalem, Israel). 2-APB was purchased from Tocris (Bristol, UK). All other chemicals were purchased from Sigma (St. Louis, MO, USA). Toxin stock solutions were made at 1000-fold concentration with distilled water or DMSO and stored at -20°C.

### Estimation of the distance between Ca^2+^ sensors and VGCCs

Extensive theoretical studies assert that buffered calcium diffusion from open calcium channels make calcium microdomain of tens of micromolar concentration (Neher, 1998). Assuming that free buffer concentration does not change very much owing to rapid diffusional replacement of Ca^2+^-bound buffer molecules with free buffer, the spatial profile of [Ca^2+^] as a function of distance from an open calcium channel (r) is given by:

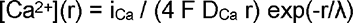

where iCa = calcium current; F = Faraday constant; DCa = diffusion constant of Ca^2+^ in cytosol.

Therefore, the ratio of [Ca^2+^] at a given distance from a Ca^2+^ channel (r) before and after the addition of BAPTA is as follows:

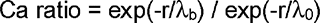

where λ0 and λb are the length constants before and after BAPTA was added, respectively. Since the length constant, λ, in the presence of free calcium buffer [B] with Ca^2+^ binding rate constant, kon, is given by

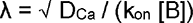

Inserting this equation into the Ca ratio equation, we have dependence of calcium ratio as a function of [BAPTA]total (x, in mM) as follows:

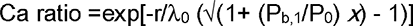

where λ0 = length constant in the presence of endogenous buffer (B0) alone; P0 and Pb,1 are buffer products (kon [B]) of B0 and 1 mM BAPTA, respectively. Finally, assuming that only the calcium influx through a calcium channel (or cluster) nearest to a vesicle is relevant to its release, and that glutamate release has the n-th power dependence on [Ca^2+^], the ratio of release (Rrls) before and after adding BAPTA has the relationship:

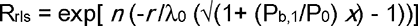

The n values for miniature and evoked EPSCs were obtained from Fig. 1D and 1F. Pb,1 is calculated as 300 mM^-1^ms^-1^ considering that kon of BAPTA = 400 mM^-1^ms^-1^, and [Ca^2+^]rest = 80 nM. Fitting the above equation to the plot of evoked or miniature EPSC frequency as a function of [BAPTA]total with setting r/λ0 and P0 as free parameters, we estimated the distance of vesicles from the calcium source (r).

### Statistical analysis

Data were are expressed as the mean ± SEM, where N represents the number of cells studied. Statistical analysis was performed using IgorPro (version 6.1, WaveMetrics, Lake Oswego, OR, USA) and OriginPro (version 9.0, OriginLab Corp., Northampton, MA, USA). Significant differences between the experimental groups were analyzed using independent or paired Student’s t-tests. P<0.05 was considered statistically significant.

---

**Supplementary Fig. 1.**
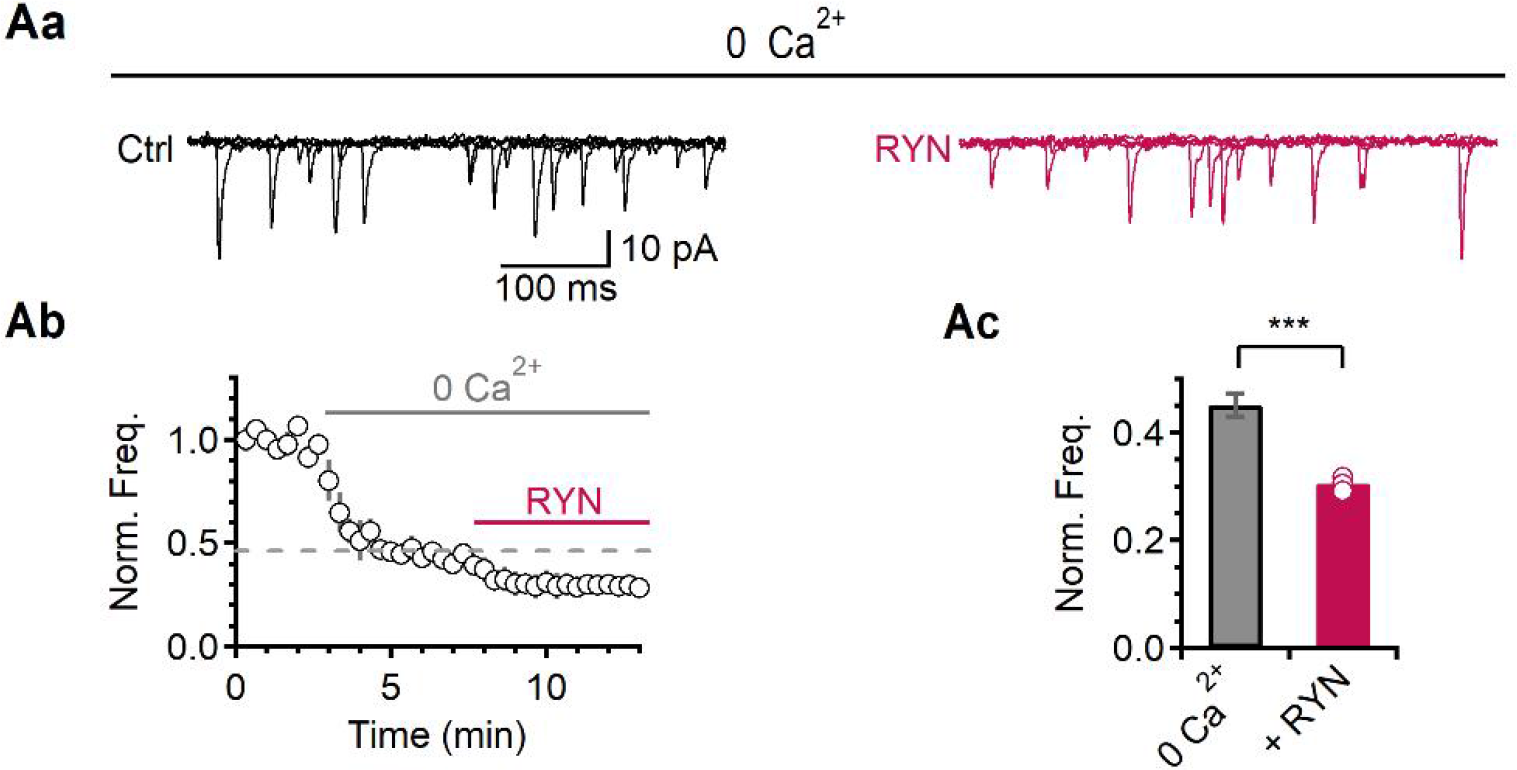
The effect of RYN on mEPSC frequency in the Ca^2+^-free extracellular solution. (Aa) Representative traces of mEPSCs in control and 5 min after applying RYN, respectively. Five 500 ms-long mEPSC traces were overlaid. (Ab) An average time course of the normalized mEPSC frequency. The solid lines indicate the presence of 0 Ca^2+^ and RYN, respectively. The data were normalized by the mean mEPSC frequency of 2 Ca^2+^. (Ac) A bar graph of average values of the normalized mEPSC frequency in RYN in the Ca^2+^-free external solution (0 Ca^2+^ *vs* RYN, 0.45 ± 0.02 *vs* 0.31 ± 0.01, compared to 2 Ca^2+^, *N* = 4). All data are represented as mean ± S.E.M., \*\*\**P*<0.001, single group mean *t*-test.

**Supplementary Fig. 2.**
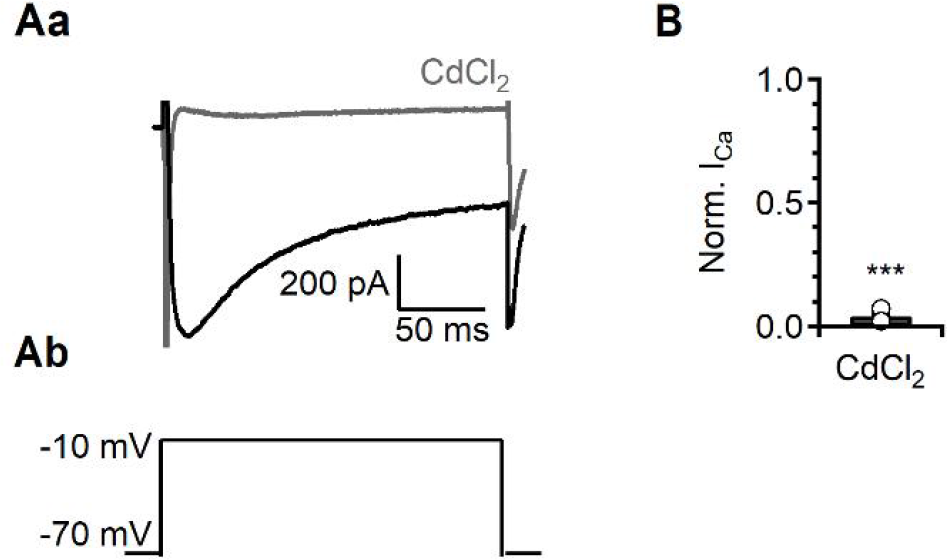
The effect of CdCl_2_ on I_Ca_. (Aa) Representative traces of I_Ca_ in control and the presence of 100 μM CdCl_2_. (Ab) The pulse protocol for recording I_Ca_. The VGCC currents were recorded by changing clamed voltage from -70 to -10 mV via step-mode in bath solution containing 25 mM tetraethylammonium (TEA), 5 mM 4-AP, 1 μM TTX, 10 μM CNQX and 100 μM PTX. (B) A bar graph of average values of the normalized I_Ca_ in CdCl_2_ (0.04 ± 0.01, *N* = 3). All data are represented as mean ± S.E.M., \*\*\**P*<0.001.

**Supplementary Fig. 3.**
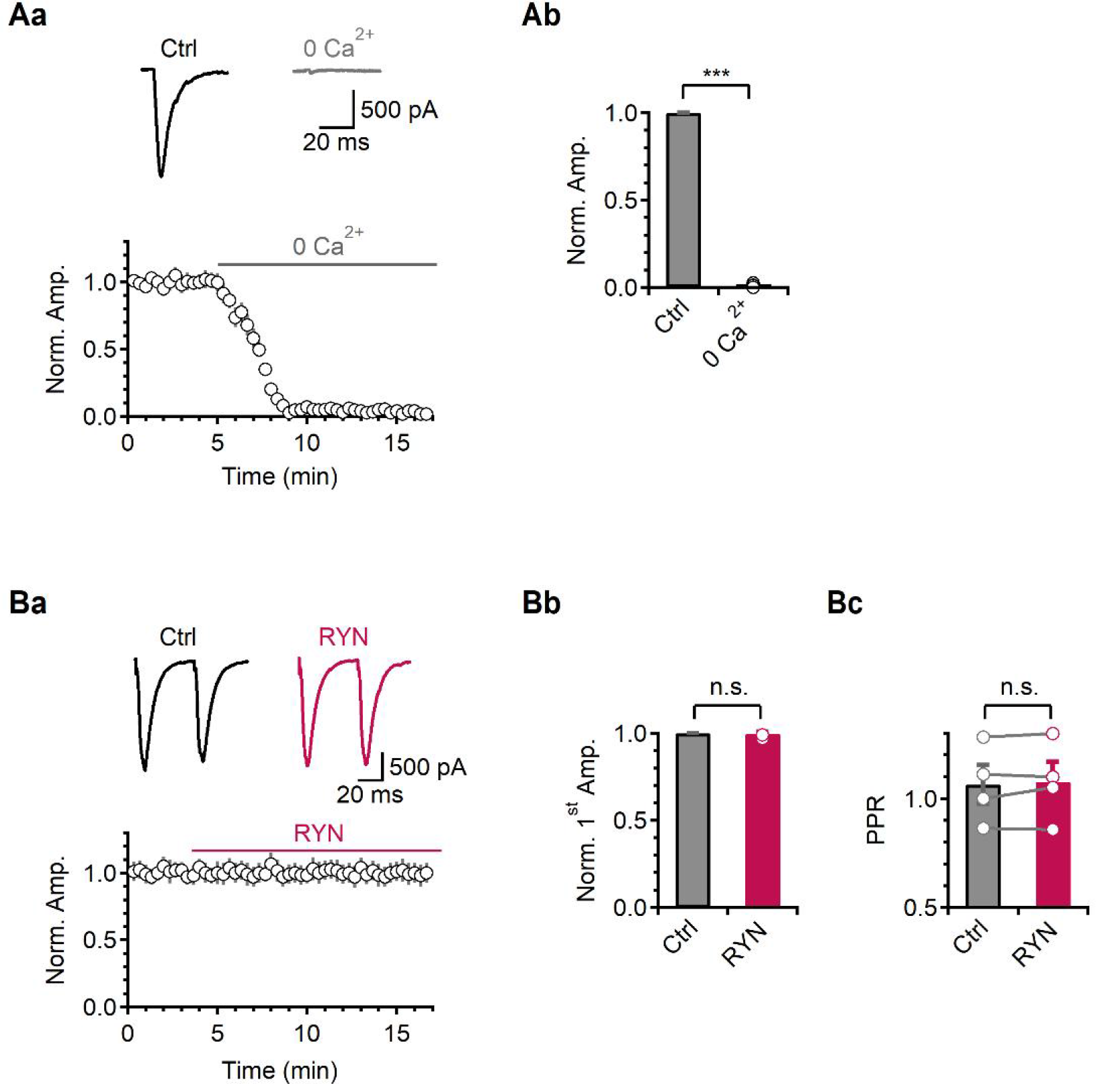
The effects of Ca^2+^ removal or RYN on eEPSC. (Aa) Top. Representative traces of eEPSC in control and 0 Ca^2+^ (grey). Bottom. An average time course of the normalized eEPSC amplitude. The solid line indicates the presence of 0 Ca^2+^. The data were normalized by the mean amplitude of the 1^st^ eEPSC amplitude of control. (Ab) A bar graph of average values of the normalized 1^st^ eEPSC amplitude (0 Ca^2+^, 0.008 ± 0.01, *N* = 7). (Ba) Top. Representative traces of eEPSC in control and 10 min after applying RYN (magenta). Bottom. An average time course of the normalized 1^st^ eEPSC amplitude. The solid line indicates the presence of RYN. The data were normalized by the mean amplitude of the 1^st^ eEPSC amplitude of control. (Bb-c) Bar graphs of average values of the normalized 1^st^ eEPSC amplitude (Bb, 0.99 ± 0.01) and PPR (Bc, Ctrl *vs* RYN, 1.06 ± 0.09 *vs* 1.07 ± 0.1; *N* = 4). All data represented as are mean ± S.E.M., \*\*\**P*<0.001, paired *t*-test; n.s. = not significant.

**Supplementary Fig. 4.**
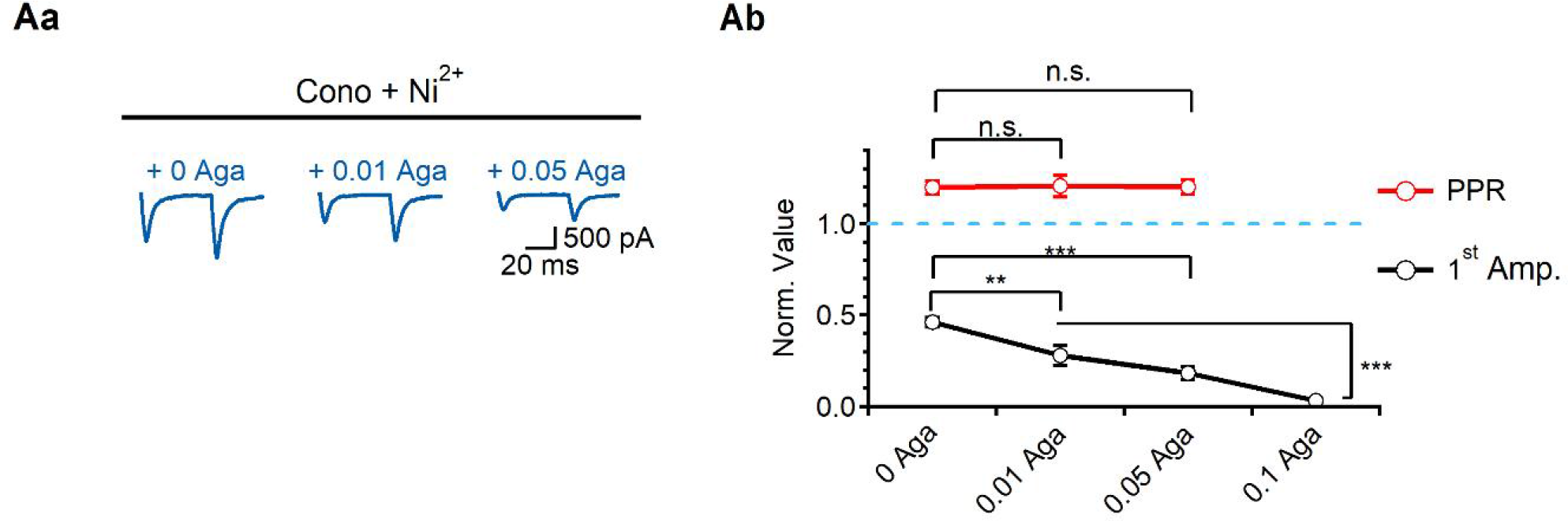
The effects of Aga on amplitudes and PPR in Cono + Ni^2+^ pre-treated cells. (Aa) Representative traces of eEPSC in Cono + Ni^2+^ pre-treated cells with dose-dependent application of Aga. (Ab) A graph of average values of the normalized 1^st^ amplitudes (black) or normalized PPR (red), compared to control (1^st^ Amp.; Cono + Ni^2+^, 0.46 ± 0.03; 0.01 μM Aga, 0.28 ± 0.05; 0.05 μM Aga, 0.18 ± 0.04; 0.1 μM Aga, 0.03 ± 0.01; PPR, Cono + Ni^2+^, 1.2 ± 0.03; 0.01 μM Aga, 1.21 ± 0.06; 0.05 μM Aga, 1.2 ± 0.04, *N* = 6). The blue dashed line indicates the value of “1”. All data are represented as mean ± S.E.M., single group mean *t* test or paired *t*-test; n.s. = not significant.

**Supplementary Fig. 5.**
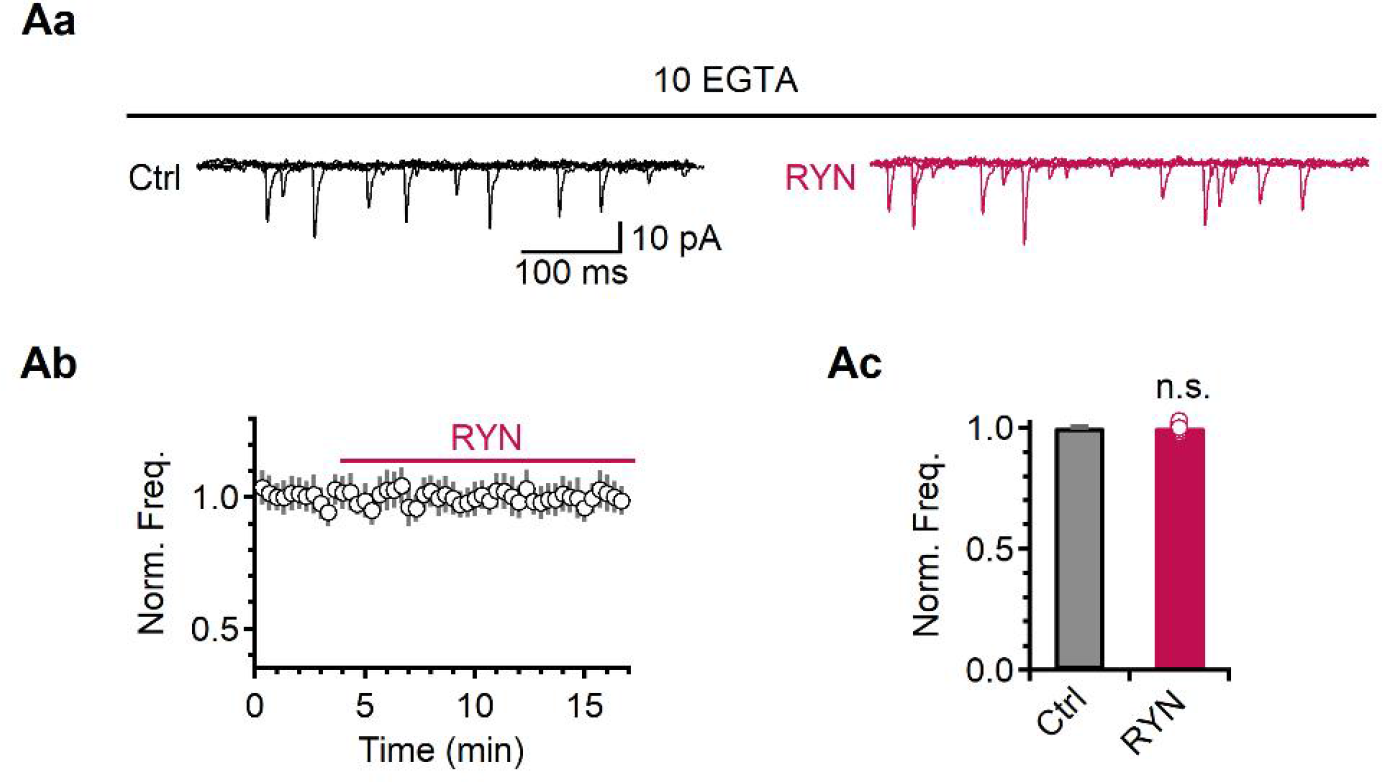
The effect of RYN on mEPSC frequency in the internal solution containing 10 mM EGTA. (Aa) Representative traces of mEPSC in control and 10 min after applying RYN. Five 500 ms-long mEPSC traces were overlaid. (Ab) An average time course of the normalized mEPSC frequency. The solid line indicates the presence of RYN. The data were normalized by the mean mEPSC frequency of control. (Ac) A bar graph of average values of the normalized mEPSC frequency in RYN (1.0 ± 0.01, compared to control, *N* = 4). All data are represented as mean ± S.E.M., single group mean *t* test; n.s. = not significant.

**Supplementary Fig. 6.**
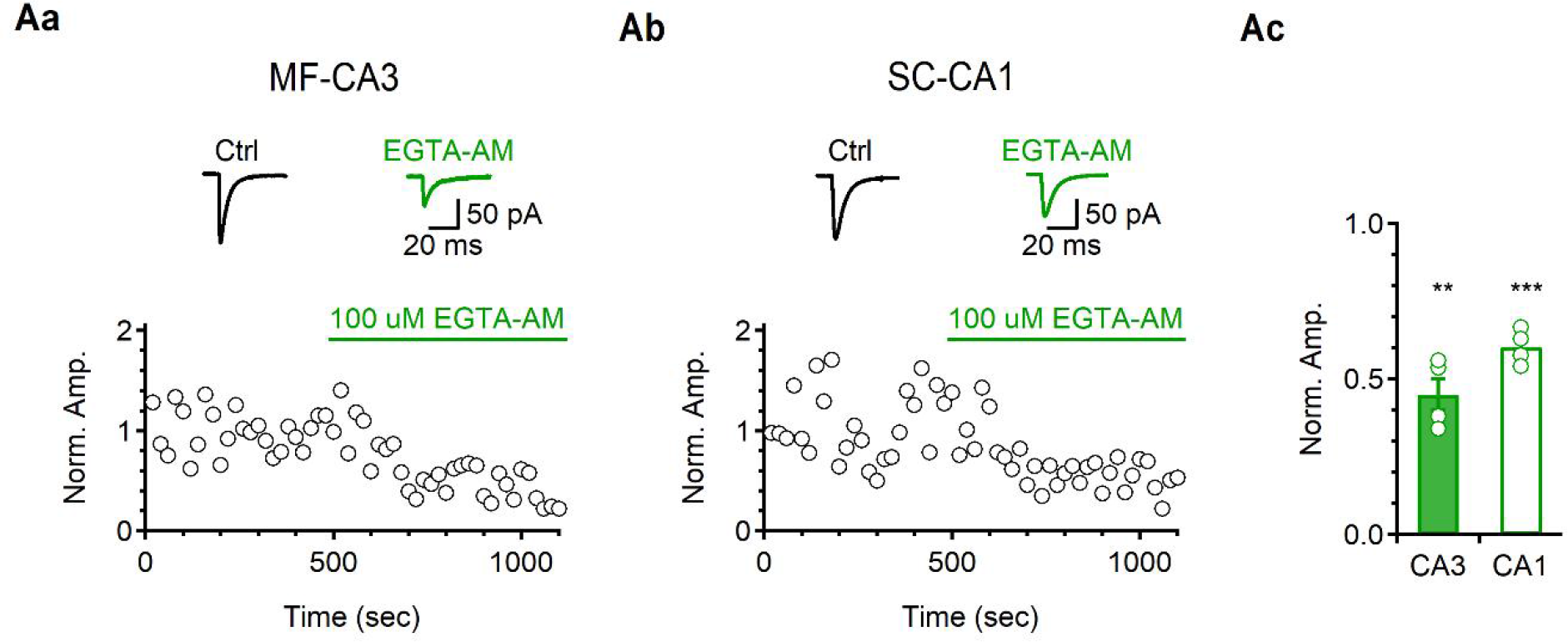
The effect of 100 μM EGTA-AM on evoked glutamate release in acute hippocampal slice. (Aa-b) Top. Representative traces of eEPSCs in control and the presence of 100 μM EGTA-AM (green) in CA3 (Aa) and CA1-PCs (Ab) in response to MF or SC stimulation, respectively. Bottom. Average time courses of the normalized eEPSC amplitude. In each time course plot, solid lines indicate the presence of 100 μM EGTA-AM. The data were normalized by the mean eEPSC amplitude of control. (Ac) A bar graph of average values of the normalized eEPSC amplitude in different condition (CA3, 0.45 ± 0.05, *N* = 4, *P* = 0.002; CA1, 0.60 ± 0.03, *N* = 4; *P* = 0.0007). All data are represented as mean ± S.E.M., single group mean *t* test; \*\**P*<0.01, \*\*\**P*<0.001, n.s. = not significant.

